# SoxB1 transcription factors are essential for initiating and maintaining the neural plate border gene expression

**DOI:** 10.1101/2023.09.28.560033

**Authors:** Elizabeth N. Schock, Joshua R. York, Austin P. Li, Ashlyn Y. Tu, Carole LaBonne

## Abstract

SoxB1 transcription factors (Sox2/3) are well known for their role in early neural fate specification in the embryo, but little is known about functional roles for SoxB1 factors in non-neural ectodermal cell types, such as the neural plate border (NPB). Using *Xenopus laevis*, we set out to determine if SoxB1 transcription factors have a regulatory function in NPB formation. Herein, we show that SoxB1 factors are necessary for NPB formation, and that prolonged SoxB1 factor activity blocks the transition from a NPB to a neural crest state. Using ChIP-seq we demonstrate that Sox3 is enriched upstream of NPB genes in early NPB cells and, surprisingly, in blastula stem cells. Depletion of SoxB1 factors in blastula stem cells results in downregulation of NPB genes. Finally, we identify Pou5f3 factors as a potential SoxB1 partners in regulating the formation of the NPB and show their combined activity is needed to maintain NPB gene expression. Together, these data identify a novel role for SoxB1 factors in the establishment and maintenance of the NPB, in part through partnership with Pou5f3 factors.

## Introduction

SoxB1 transcription factors (Sox1/2/3) are well known for their roles during early embryonic development in establishing and maintaining pluripotency and promoting neural progenitor cells (Avilion et al., 2003; Kishi et al., 2000; Zhang and Cui, 2014). Prior to gastrulation, SoxB1 factors are expressed in cells of the embryo proper in vertebrate and invertebrate chordate species (Avilion et al., 2003; Buitrago-Delgado et al., 2018; Cattell et al., 2012; Okuda et al., 2006; Rex et al., 1997). Homozygous Sox2 murine mutants fail to develop past blastocyst stages, highlighting the importance of these transcription factors in cell survival during early embryonic development (Avilion et al., 2003). As gastrulation and lineage restriction commence, SoxB1 expression becomes restricted to the ectoderm and eventually to neuroectodermal cells prior to the onset of neurulation (Avilion et al., 2003; Buitrago-Delgado et al., 2018; Rex et al., 1997). During this period, the ectoderm is being patterned into three regions: neural plate, neural plate border, and non-neural ectoderm (Sasai and De Robertis, 1997). While SoxB1 factor expression ultimately becomes restricted to neural cells where they function redundantly to maintain a neural progenitor state (Bylund et al., 2003), SoxB1 factors are also expressed in non-neural ectodermal lineages. Lineage tracing experiments in chick have shown that initially Sox2-positive cells contribute not only to the neural plate, but also the epidermis and neural plate border (Roellig et al., 2017), but whether SoxB1 factors play functional roles in establishing these non-neural ectodermal cell types remains unclear.

During gastrulation, neural plate border cells arise lateral to neural plate cells and give rise to two other cell types, the neural crest and pre-placodal ectoderm (Groves and LaBonne, 2014; Pla and Monsoro-Burq, 2018). BMP, Wnt, and FGF signaling pathways have all been implicated in the formation of the neural plate border and promote the expression of neural plate border genes including *pax3/7*, *zic1,* and *msx1* (Garnett et al., 2012; Marchal et al., 2009; Tribulo et al., 2003). These transcription factors, in turn, help to promote/stabilize expression of each other and to activate neural crest or placodal gene expression (Hong and Saint-Jeannet, 2007; Monsoro-Burq et al., 2005; Sato et al., 2005). While functional roles for neural plate border-specific transcription factors are defined, less is known about transcriptional inputs outside of signaling cascades that contribute to the early establishment of this cell type.

SoxB1 factors are promising candidate transcription factors for regulating early neural plate border gene expression. Although SoxB1 factors are well known regulators of neural progenitor cells (Papanayotou et al., 2008), they are expressed in early neural plate border cells and recent genomic analysis has observed Sox2 binding in neural plate border/early neural crest cells (Hovland et al., 2022; Roellig et al., 2017). These Sox2 enriched regions of chromatin close as cells transition from a neural plate border to a definitive neural crest state (Hovland et al., 2022). Additionally, many neural plate border factors, including *zic1* and *pax3/7*, also have functional roles in neural cell types, suggesting overlap in the transcriptional regulation of these cells (Aruga et al., 2002a; Aruga et al., 2002b; Lin et al., 2016). Given these findings, we hypothesized that SoxB1 factors may promote neural plate border gene expression during gastrulation.

Here, we investigate a role for SoxB1 factors in the formation of the neural plate border using *Xenopus laevis*. We find that SoxB1 factor expression overlaps with the forming neural plate border until neurulation and is lost in those cells as definitive neural crest cells form. Using morpholino-mediated knockdown we show that SoxB1 factors are necessary for neural plate border formation. Forced SoxB1 factor expression promotes and prolongs a neural plate border state, delaying the formation of neural crest cells. Through ChIP-seq analyses, we find that Sox3 directly regulates expression of core neural plate border genes, and this transcriptional regulation begins in blastula stem cells. Finally, we provide evidence that SoxB1 factors partner with Pou5f3 factors to promote neural plate border formation. Together, these results identify a key functional role for SoxB1 factors in the formation of the neural plate border.

## Results

### Sox3 co-localizes with neural plate border marker pax3 through gastrulation

A role for SoxB1 factors in establishing the neural plate border would require they be expressed in that region during gastrulation. We therefore examined whether SoxB1 protein co-localizes with the key neural plate border factor *pax3*. As Sox2 and Sox3 are functionally redundant (Bylund et al., 2003) and have identical expression patterns at early stages of development (Sup. Fig. 1A) we focused on Sox3 localization as its expression is significantly higher than Sox2 at these stages (Sup. Fig 1B). A *Xenopus*-specific Sox3 antibody (Horr et al., 2023) was used in combination with a hybridization chain reaction (HCR) probe for *pax3* to mark the forming neural plate border (Choi et al., 2018). We asked if *pax3*-positive neural plate border cells were also Sox3-postive during gastrulation and neurulation. We observed that during gastrulation (stage 11.5) *pax3* expressing cells are Sox3-postive (Fig. 1A). Co-expression of Sox3 and *pax3* appears to decrease laterally by neural plate stages (stage 13) with little to no overlap by mid-neurula stages (stage 15) when definitive neural crest cells are present (Fig. 1A).

**Figure 1.**
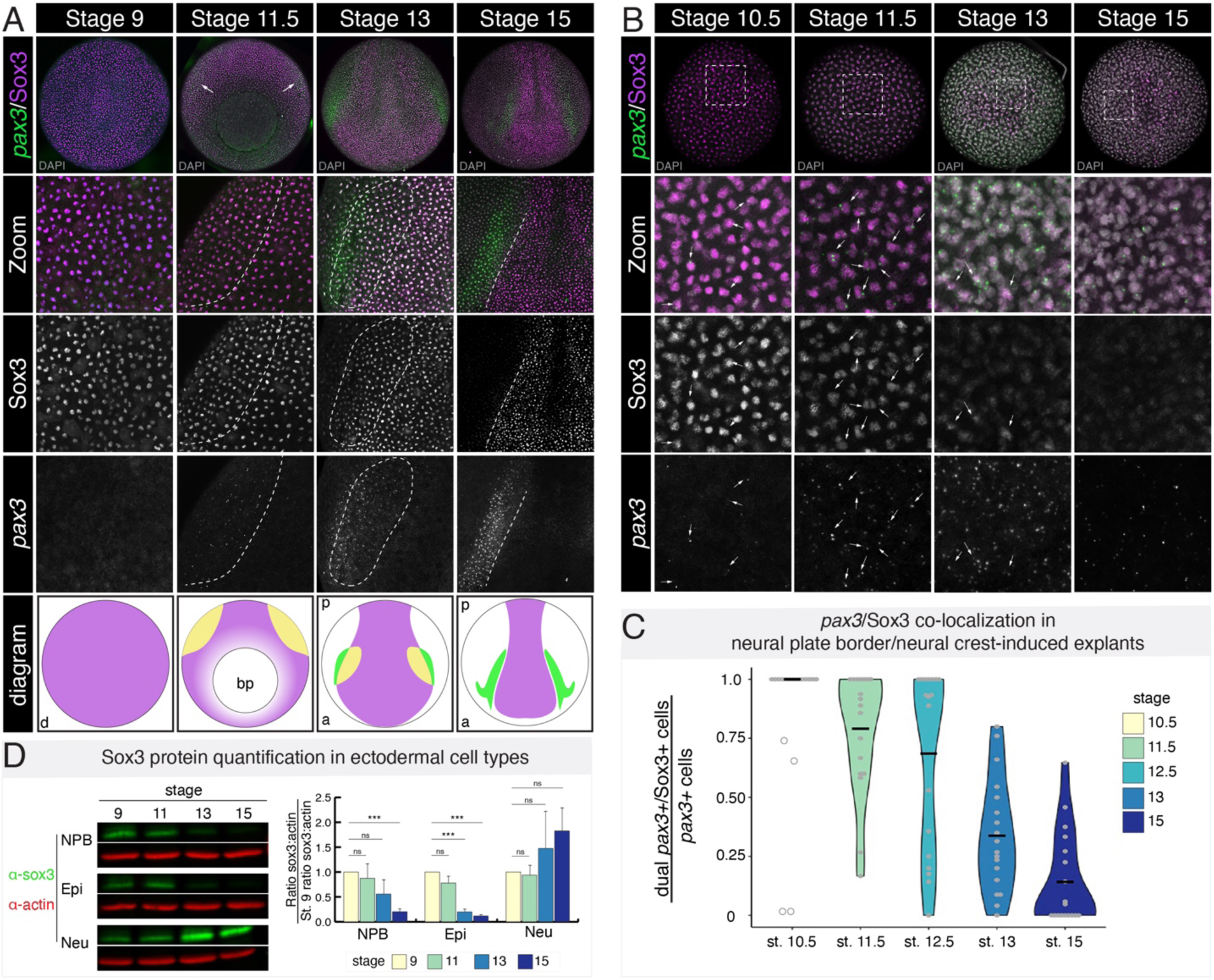
Sox3 protein co-localization with *pax3* transcripts during gastrulation and co-localization decreases with the onset of neurulation. (A) Whole embryos immunostained for Sox3 (magenta) and probed with HCR oligos for *pax3* (green) across multiple developmental stages (blastula through late neurula). DAPI is shown in gray. Diagrams show domains of co-localization (yellow) and at each time point. Sox3 protein localization is displayed in magenta and *pax3* transcripts in green in diagrams. (B) Neural plate border/neural crest-induced explants immunostained for Sox3 (magenta) and probed with HCR oligos for intronic *pax3* (green), marking nascent transcripts, across multiple developmental stages (blastula through late neurula). DAPI is shown in gray. Arrows denote cells where nascent *pax3* transcripts co-localize with nuclear Sox3 protein. (C) Quantification of co-localization, as shown by violin plots, between Sox3 protein and nascent *pax3* transcripts over developmental time. Outlier data points are displayed as hollow circles. Means are displayed as black horizonal bars. (D) Western blot for endogenous Sox3 protein (green) and actin (red) in neural plate border-induced explants, epidermal explants, and neural-induced explants at stages 9, 11, 13, and 15. Bar graph shows the quantification of western signal where the ratio of Sox3 to actin is normalized to stage 9. Error bars show standard deviation. ns=not significant; (***) p<0.001; dorsal (d); anterior (a); posterior (p); blastopore (bp); neural plate border (NPB); epidermis (Epi); neural (Neu)

To more precisely quantify the degree of cellular co-localization between Sox3 and *pax3*, we used a blastula stem cell explant system where dissected blastula stem cells can be induced to form any cell type in the embryo given the appropriate developmental cues (Ariizumi and Asashima, 2001). To pharmacologically induce a neural plate border state, we treated dissected explants with the small molecules CHIR and K02288 (BMPi), a Wnt agonist and BMP antagonist, respectively (Huber and LaBonne, in revision). Neural plate border-induced explants were collected at time points corresponding with early gastrulation (stage 10.5) through mid-neurulation (stage 15) and immunostained for endogenous Sox3 and probed for nascent *pax3* transcripts, though use of an intronic probe set (Fig. 1B; Sup. Fig. 2A). By examining nascent transcripts, we can determine if Sox3 protein is present in the nuclei of cells that are actively synthesizing *pax3* transcripts. Our co-localization analysis revealed that at early gastrula stages (stage 10.5) 100% of nuclei with nascent *pax3* transcripts are Sox3-postive (Fig. 1C). We observed that *pax3* expression and nuclear Sox3 immunoreactivity remained highly correlated (79% co-localization (stage 11.5); 69% co-localization (stage 12.5)) until the start of neurulation (stage 13), when a decrease in co-localization (34%) was observed (Fig. 1C). By stage 15, only 14% of the *pax3*-positive cells were also Sox3-positive (Fig. 1C). Nascent *pax3* transcripts were infrequently observed in control epidermal explants (Sup. Fig. 2B-D). This analysis indicates that Sox3 protein is present in newly formed *pax3*-positive neural plate border cells during early to mid-gastrulation before decreasing at neurula stages as definitive neural crest genes become expressed (Sup. Fig. 2E).

We also examined changes in levels of Sox3 protein over time in three different ectodermal cell populations: neural plate border, epidermal, and neural cells via western blot. Untreated blastula stem cells become epidermis due to endogenous BMP signaling (Wilson et al., 1997) and a neural state can be induced using a high dosage of BMPi (Johnson et al., 2022). Explants for each ectodermal cell type were collected at time points corresponding with blastula stage (stage 9) through mid-neurula (stage 15) and endogenous levels of Sox3 were assessed via western blot and quantified, normalizing using actin (Fig. 1D). Consistent with what we observe by immunofluorescence, the levels of Sox3 protein were highest in neural plate border cells during gastrulation (stage 11), with a significant decrease (80% decrease from stage 9; p=0.001) in Sox3 protein by mid-neurulation (stage 15). In contrast, Sox3 protein increased in neural-induced explants over time (Fig. 1D). Sox3 was also present at high levels in epidermal cells at stage 11 but decreased significantly (80% decrease from stage 9; p=0.001) by the start of neurulation (stage 13). Together with our immunofluorescence data, these findings demonstrate that at the onset of neural plate border formation nuclear Sox3 protein is present in these cells, but co-expression decreases during neurulation.

### SoxB1 transcription factors are necessary for neural plate border formation

The presence of Sox3 in forming neural plate border cells during gastrulation is consistent with a potential role for SoxB1 factors in regulating their formation. To determine if SoxB1 factors are necessary for neural plate border gene expression, we designed and validated translation-blocking morpholinos targeting both alloalleles of *sox2* and *sox3* (Sup. Fig. 3). We next performed double morpholino-mediated knockdown of Sox2 and Sox3 and examined neural plate border formation via *in situ* hybridization. We observed a loss of neural plate border markers *pax3*, *zic1*, and *msx1* whereas expression of *tfap2a* remained unchanged (Fig. 2A; Sup. Fig. 4A). We found that neural plate border gene expression could be rescued by expressing with a single SoxB1 factor (*sox3*) (Fig. 2B). We also examined the effects of *soxb1* depletion in blastula stem cells induced to a neural plate border state. We observed that neural plate border genes *pax3* and *msx1* were poorly/weakly induced in *soxb1* morphants compared to controls (Fig. 2C; Sup. Fig. 4B). We also found that expression of several additional neural plate border genes (*dlx5*, *prdm1*, *klf17*) was significantly reduced in double morphant neural plate border-induced explants via qPCR (Fig. 2D) Together, these data indicate that SoxB1 factors are required for neural plate border formation.

**Figure 2.**
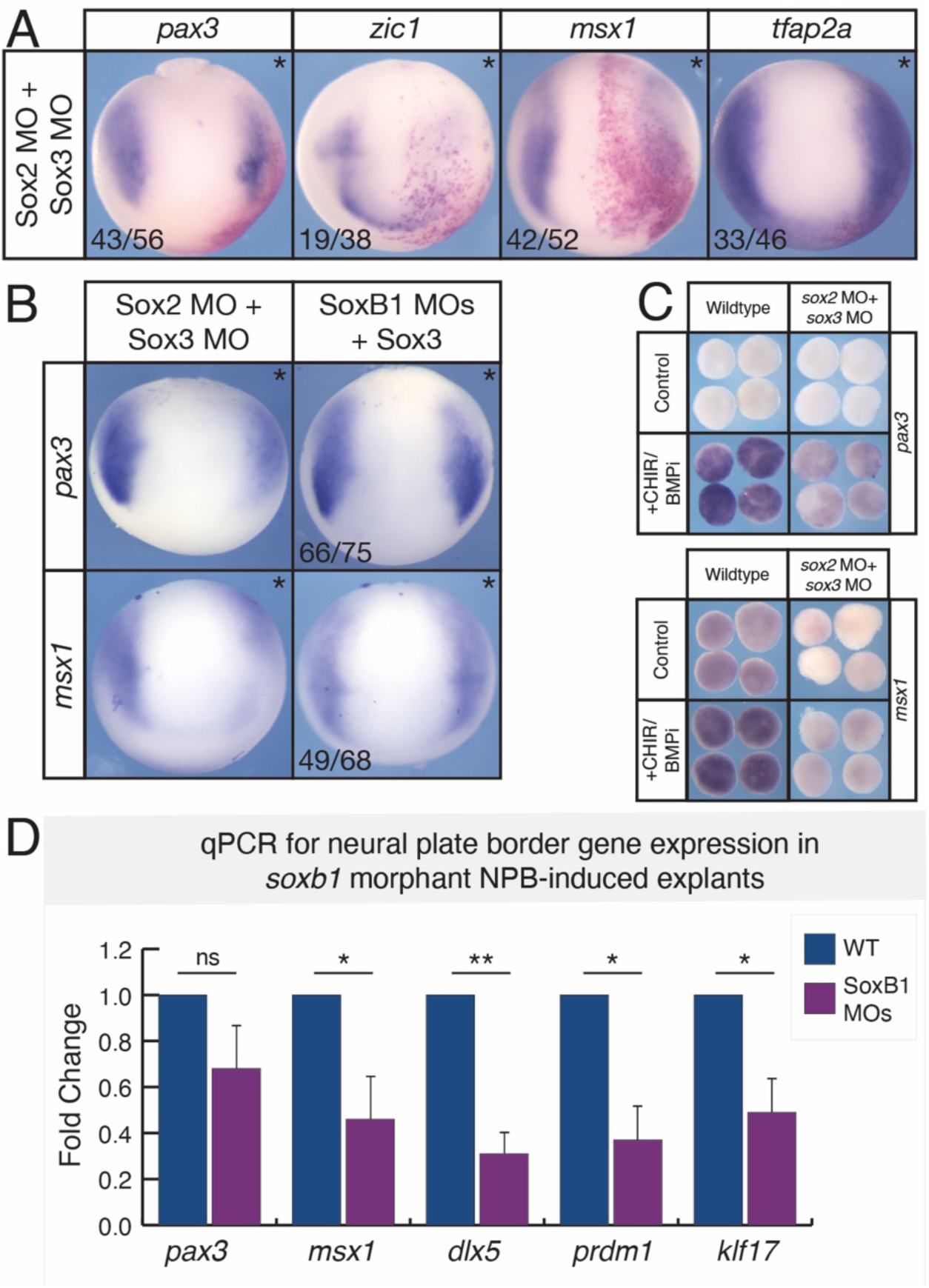
SoxB1 factors are necessary for neural plate border formation. (A) *In situ* hybridization for *pax3*, *zic1, msx1*, and *tfap2a* in stage 12.5-13 embryos unilaterally injected with *sox2* and *sox3* morpholino (*denotes injected side). Beta-galactosidase (red) was used as a lineage tracer. (B) *In situ* hybridization for *pax3* and msx1 in stage 12.5-13 embryos unilaterally injected with *sox2* and *sox3* morpholinos or rescued embryos which were co-injected with *sox3* mRNA (*denotes injected side). Fluorescein dextran was used as a lineage tracer and embryos were presorted for left/right side targeting. (C) *In situ* hybridization for *pax3* and *msx1* on wildtype and *soxb1* morphant neural plate border-induced explants (stage 12.5). (D) qPCR examining gene expression fold changes in neural plate border-induced explants (stage 12.5) comparing wildtype neural plate border-induced cells (blue) to *soxb1* morphant neural plate border-induced cells (magenta). Error bars show standard deviation. (*) p<0.05; (**) p< 0.01; (***) p<0.001; morpholino (MO); neural plate border (NPB)

### SoxB1 factor expression blocks the transition from a neural plate border to a neural crest state

To determine if SoxB1 factor activity promotes neural plate border gene expression we performed gain-of-function experiments. mRNA encoding either *sox2* or *sox3* was injected into one blastomere at the two or four cell stage and embryos were assessed for changes in gene expression via *in situ* hybridization. Expression of either *sox2* or *sox3* resulted in expanded expression of *pax3* and *zic1* in early neurula embryos (stage 13; Fig. 3A). We next wanted to determine if prolonged SoxB1 activity extended neural plate border gene expression in the embryo. Our western blot and imaging analyses indicated that Sox3 protein is largely absent in neural crest cells by the middle of neurulation (stage 15), so we examined neural plate border gene expression at this stage and later. We found that expression of both *pax3* and *zic1* was expanded into the presumptive neural crest domain in mid-neurula embryos (Fig. 3B, arrowhead). We next examined neural crest gene expression at this stage and observed a loss of neural crest as evidenced by loss of *foxd3* and *snai2* gene expression (Fig. 3C; Sup. Fig. 5A) (Buitrago-Delgado et al., 2018; Rogers et al., 2009; Wakamatsu et al., 2004). Supporting these results, when we examined expression of *foxD3* and *pax3* in *sox3* expressing embryos using HCR, we found that *pax3*-positive cells were present in the domain where *foxd3* expression had been lost (Fig. 3D, arrowhead). We also examined markers for other ectodermal lineages (epidermis, placode, and neural) in *soxb1* expressing embryos, and found that expression of those lineage markers was also decreased (Sup. Fig. 5B). These data suggest that SoxB1 activity causes cells that would normally become definitive neural crest to instead be retained in neural plate border state.

**Figure 3.**
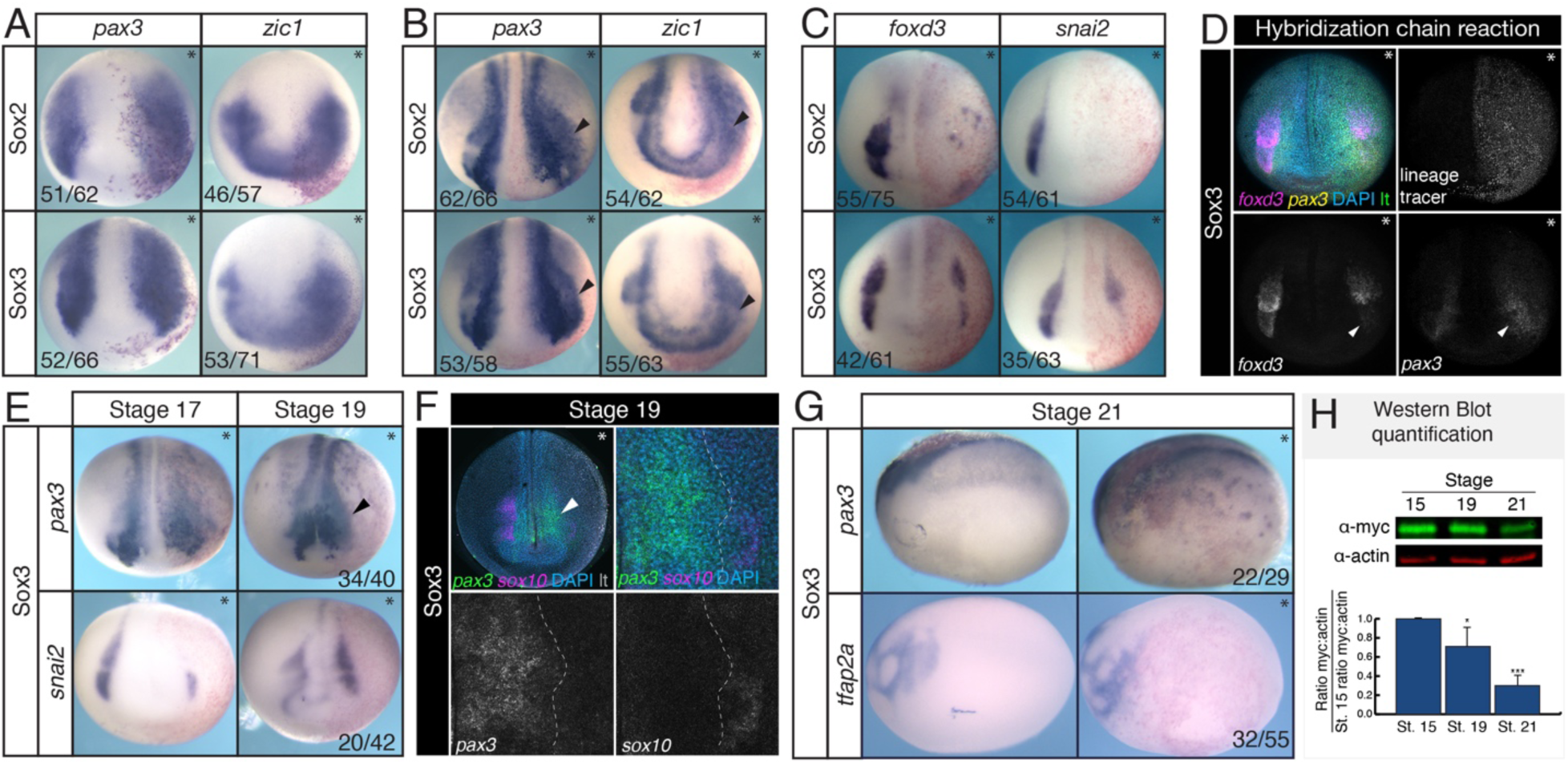
SoxB1 expression blocks the transition from neural plate border to neural crest gene expression. (A-C) *In situ* hybridization in embryos unilaterally expressing *sox2* or *sox3* mRNA (* denotes injected side). Beta-galactosidase (red) was used as a lineage tracer. (A) *pax3* and *zic1* in early neurula embryos; (B) *pax3* and *zic1* in late neurula embryos, arrowhead denotes expanded domain of expression; (C) *foxd3* and *snai2* in late neurula embryos. (D) Hybridization chain reaction for *foxd3* (magenta) and *pax3* (yellow) in embryos unilaterally expressing *sox3* mRNA (* denotes injected side). Myc antibody staining (green) was used as a lineage tracer. Arrowhead denotes region with loss of *foxd3* expression and expanded *pax3* expression. (E) *pax3* and *snai2* in stage 17 and stage 19 embryos unilaterally expressing *sox3* mRNA. Arrowhead denotes expanded domain of expression. (F) Hybridization chain reaction for *sox10* (magenta) and *pax3* (green) in embryos unilaterally expressing *sox3* mRNA (* denotes injected side). Myc antibody staining (gray) was used as a lineage tracer. (G) *In situ* hybridization for *pax3* and *tfap2a* in stage 21 in embryos unilaterally expressing *sox3* mRNA. (H) Time course western blot with quantification for myc-tagged Sox3 protein levels in the embryo where the ratio of myc to actin is normalized to stage 15. (*) p<0.05; (***) p<0.001; lineage tracer (lt)

If prolonged SoxB1 activity prevents the transition from a neural plate border to a neural crest state, it is possible that the neural crest may recover later in development once SoxB1 proteins turn over, similar to what we observe during neurulation in wildtype embryos (Fig. 1). To test this, we performed *in situ* hybridization for *pax3* and *snai2* on *sox3* injected embryos collected at stage 19. We found that *pax3* expression continues to be laterally expanded at this stage while *snai2* expression either remains lost or has partially recovered, but displays delayed migration (Fig. 3E; Sup. Fig. 5C). We also examined localization of *pax3* and *sox10* expressing cells in *sox3* injected embryos using HCR. We observed that sox*10* expression is absent in the domain of expanded *pax3* expression (Fig. 3F). By stage 21, wildtype neural crest cells have delaminated and begun to migrate (Theveneau and Mayor, 2012). When we examined expression of *pax3* and *tfap2a*, a marker for migrating neural crest cells, we found that anterior *pax3* expression is medially restricted and *tfap2a* expressing cells are present, also indicating a recovery of the neural crest (Fig. 3G; Sup. Fig. 5C). Western blot analysis indicated that myc-tagged Sox3 protein levels were reduced by 30% (p=0.04) at stage 19 compared and 70% (p=1.2×10^-4^) at stage 21 relative to levels at stage 15 (Fig. 3H). These data indicate that transitioning from a neural plate border to neural crest state is correlated with decreased levels of SoxB1 protein.

### Sox3 is enriched upstream of neural plate border genes

To determine if Sox3 directly regulates neural plate border gene expression we performed ChIP-seq in mid-gastrula (stage 11.5) explants expressing myc-tagged *sox3* that had been induced to a neural plate border state with *wnt8a* and *chordin* mRNA (LaBonne and Bronner-Fraser, 1998) (Fig. 4A). We validated that these *sox3* injected explants were bona fide neural plate border cells by assaying for *pax3* expression (Sup. Fig. 6). Peak calling identified 5,300 peaks enriched for Sox3 binding. When we examined genomic regions of known neural plate border genes (*pax3*, *zic1*, *msx1*, *dlx6*, *klf17*, *prdm1*) we observed Sox3 enrichment upstream (∼5 kb or less), or in putative regulatory regions of introns of these genes (Fig. 4B).

**Figure 4.**
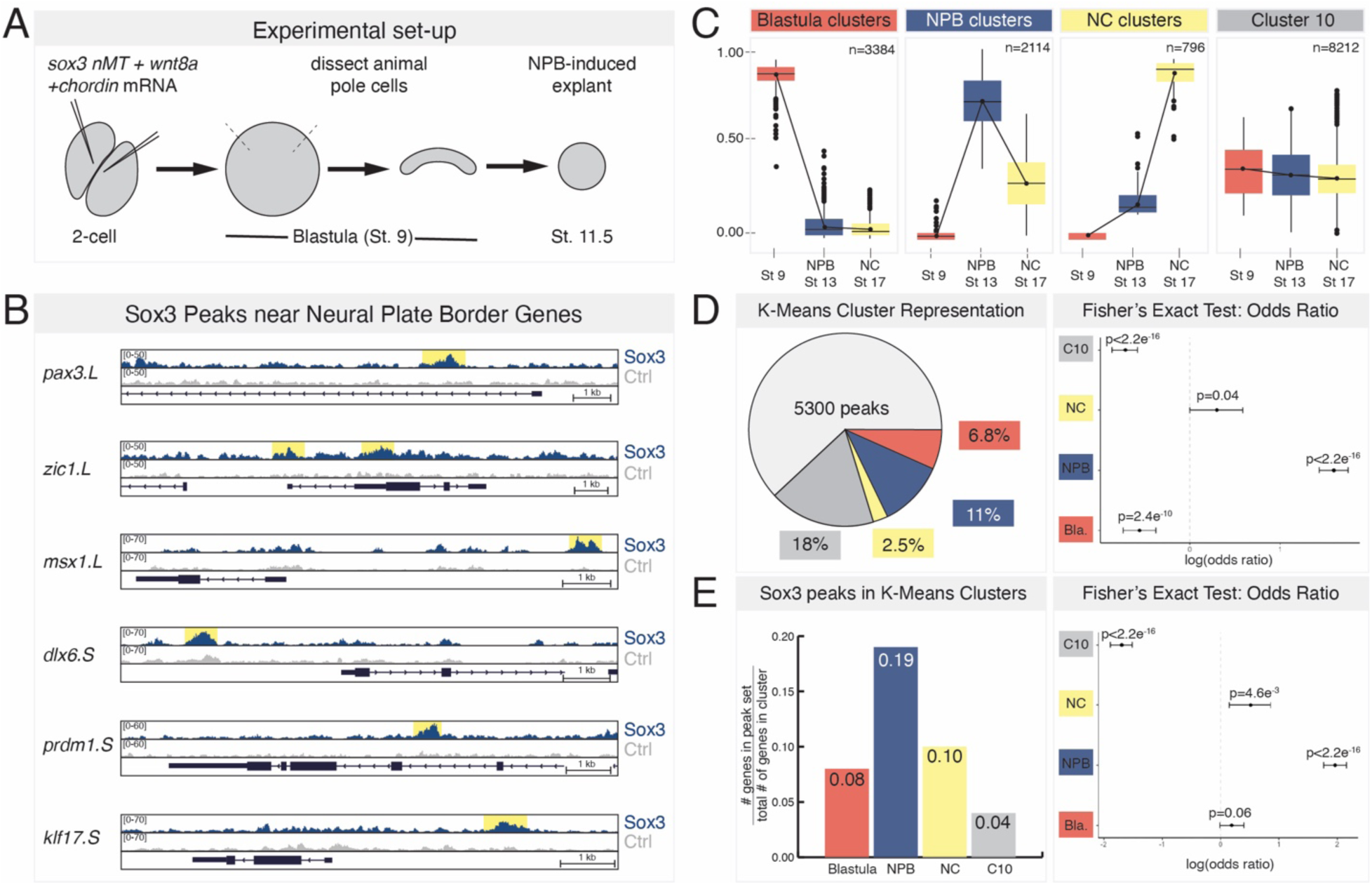
ChIP-seq in early neural plate border cells reveals Sox3 enrichment at multiple neural plate border genes. (A) Schematic of experimental design for Sox3 ChIP-seq experiments in neural plate border-induced explants. (B) Genome browser view of Sox3 peaks at loci for neural plate border genes (*pax3*, *zic1*, *msx1*, *dlx6*, *prdm1*, and *klf17*). (C) Representative k-means clusters for transcriptome data for blastula stem cells (stage 9), neural plate border-induced explants (stage 13), and neural crest-induced explants (stage 17) (Adapted from York et al., in revision). (D) Percentage of Sox3 peaks (5300 peaks total) associated with k-means cluster genes and accompanying forest plot for odds ratio (E) Percentage of genes in k-means clusters with associated Sox3 ChIP peaks and accompanying forest plot for odds ratio. Blastula cluster (red); neural plate border (NPB) cluster (blue); neural crest (NC) cluster (yellow); Cluster 10 (C10; gray)

We next wanted to determine if determine if Sox3 enrichment occurs more frequently at neural plate border genes than non-neural plate border genes. To do so, we utilized existing k-means analysis of transcriptome data from *Xenopus* blastula stem cells (stage 9) and neural plate border/neural crest-induced explants (stage 13/17) (York et al., in revision; Godichon-Baggioni et al., 2019; Rau and Maugis-Rabusseau, 2018). Specifically, we utilized three clusters that display distinct signatures for blastula (*pou5f3.3*, *foxi2*), neural plate border (*pax3*, *zic1*), and neural crest (*foxd3*, *snai2*) genes, and a fourth cluster, cluster 10 (C10), consisting of genes lacking dynamic expression changes that includes many housekeeping genes (*cox2*, *gapdh*, and *ubc* (Silver et al., 2008)) (Fig. 4C). We used these gene lists to assess if Sox3 occupancy was more highly enriched in neural plate border genes (blue) than non-neural plate border genes.

We compared the list of genes associated with our Sox3 ChIP peaks, as defined by the nearest TSS, to these different k-means clusters and found that, of the 5,300 peaks, 11% were associated with genes (n=596) in the neural plate border cluster (Fig. 4D). We calculated an odds ratio for this value using a Fisher’s exact test and found this association to be highly significant (odds ratio=3.02; p<2.2×10^-16^). In contrast, the other three cluster were not significantly enriched for Sox3, as determined by Fisher’s exact testing (Fig. 4D). We also performed the reciprocal analysis and asked how many genes in each cluster group have an associated Sox3 ChIP peak. We found that of the 2,114 genes in the neural plate border cluster, 19% have a corresponding Sox3 peak (n=400). In contrast, Sox3 peaks only represented 10% (NC), 8% (blastula), and 4% (C10) of the genes in the other three clusters (Fig. 3E). We again performed Fisher’s exact tests on these data and observed a high odds ratio (odds ratio=3.39; p<2.2×10^-16^) for the neural plate border cluster, suggesting that Sox3 is more highly enriched at neural plate border genes than at non-neural plate border genes.

Finally, we performed DeSeq2 on existing transcriptome data from stage 13 epidermal versus neural plate border-induced explants (York et al., in revision) and identified genes with differential expression (p_adj_<0.05, log FC>1.5) in both cell types (Sup. Fig. 7). Of the 1,675 differentially expressed genes in neural plate border cells, 452 were found to have an associated Sox3 ChIP peak (odds ratio= 3.88; p<2.2×10^-16^) (Sup. Fig. 7). In contrast, Sox3 was not enriched at differentially expressed epidermal genes (odds ratio=0.74; p=4.2×10^-5^). Overall, this ChIP-seq data analysis indicates that Sox3 is enriched at regulatory regions for neural plate border genes.

### Sox3 regulates neural plate border gene expression in blastula stem cells

Previous work from our lab has shown that a number of neural plate border genes are initially expressed in blastula stem cells (Buitrago-Delgado et al., 2015; York et al., in revision). Given that SoxB1 factors are robustly expressed in blastula stem cells, we next wanted to ask if SoxB1 factors may be regulating expression of neural plate border genes at blastula stages. To determine if this was the case, we performed ChIP-seq in blastula stem cells expressing myc-tagged *sox3* mRNA, at near-endogenous protein levels (Sup. Fig. 8). Peak calling identified ∼124,000 Sox3 peaks. As expected, we found Sox3 enriched upstream of many pluripotency genes, including *pou5f3* factors (Oct3/4), *ventx2.2* (functional equivalent of nanog), and *lin28a* (Fig. 5A). Sox3 enrichment was also observed near neural plate border genes, and we found that these peaks were highly correlated with the Sox3 peaks identified in our neural plate border ChIP data set, occupying the same genomic loci (Fig. 5B). Interestingly, we also observed Sox3 enrichment at blastula stages near genes for other ectodermal cell types: neural, epidermal, placodal, and neural crest (Fig. 5C); however, Sox3 occupancy was not maintained for these genes in neural plate border cells (Fig. 5D). These data suggest that SoxB1 factors, which are known to function as pioneer factors (Soufi et al., 2015), bind to genes associated with many lineages in blastula stem cells, but by mid-gastrula stages Sox3 occupancy is only retained at neural plate border genes.

**Figure 5.**
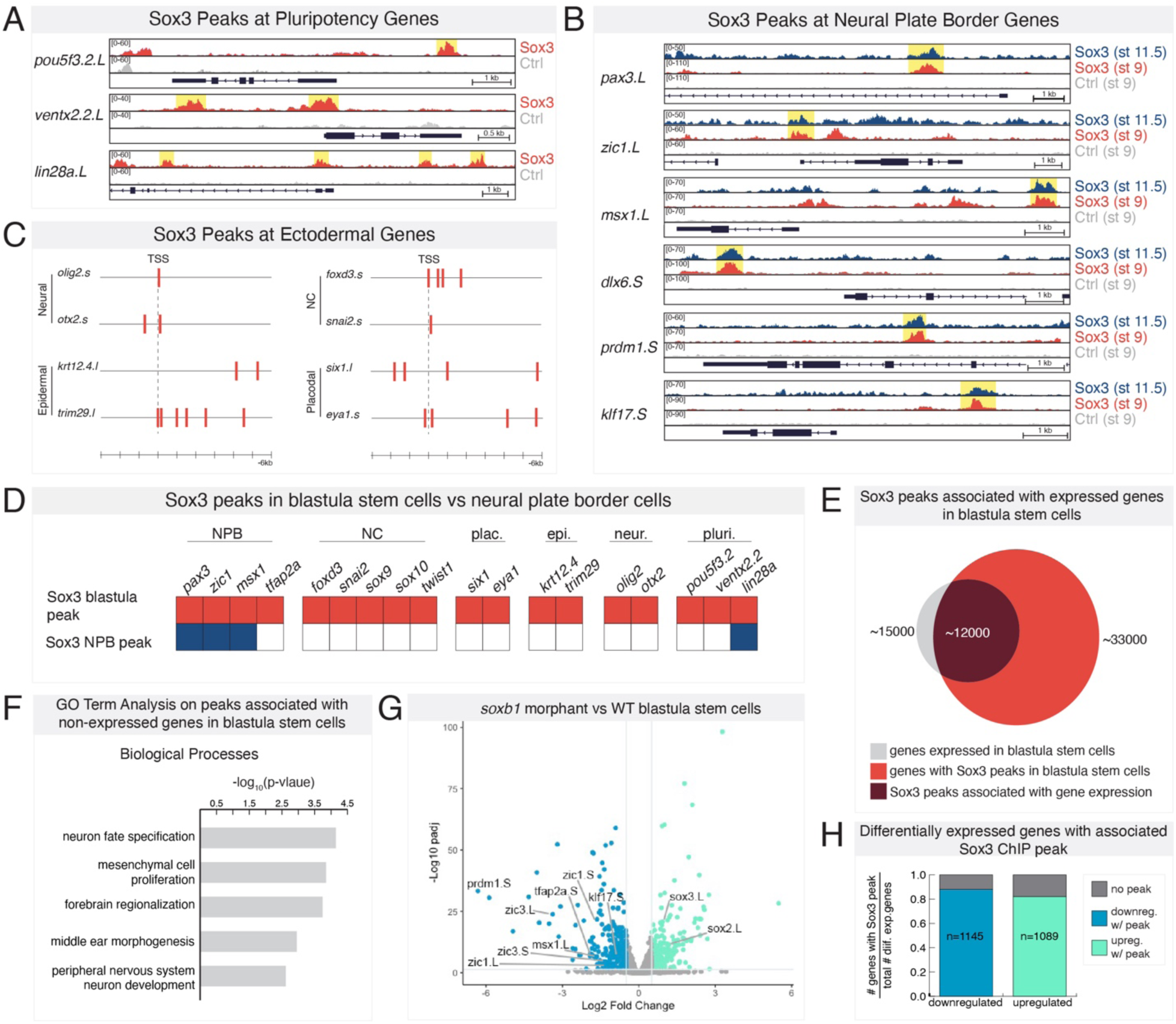
Sox3 is enriched upstream of neural plate border and other ectodermal genes in blastula stem cells. (A) Genome browser view of Sox3 ChIP peaks at loci for pluripotency genes (*pou5f3.2*, *ventx2.2*, and *lin28a*). (B) Genome browser view of Sox3 ChIP peaks at loci for neural plate border genes (*pax3*, *zic1*, *msx1*, *dlx6*, *prdm1*, and *klf17*) in blastula stem cells (red) and neural plate border cells (blue). (C) Diagram of Sox3 peaks (red bars) near the transcription start site (TSS) of neural (*olig2*, *otx2*), epidermal (*krt12.4*, *trim29*), neural crest (NC; *foxd3*, *snai2*), and placodal genes (*six1*, *eya1*). (D) Graphical summary of genome occupancy in blastula stem cells and neural plate border cells with colored boxes (red/blue) indicating Sox3 enrichment. (E) Venn diagram showing overlap (maroon) between blastula stem cell transcriptome (gray) and Sox3 peaks in blastula stem cells (red). (F) GO term analysis for genes associated with Sox3 peaks that are not expressed in blastula stem cells. (G) Volcano plot for differentially expressed genes between wildtype blastula stem cells and *soxb1* morphant blastula stem cells. Downregulated genes are shown in blue and include several neural plate border genes. Upregulated genes are shown in green. (H) Percent of differentially expressed genes with an associated blastula stage Sox3 peak. Neural plate border (NPB); neural crest (NC)

Additionally, we noted that some Sox3 peaks (stage 9) were associated with genes that are not expressed at blastula stages (ex: *pax3, olig2, six1*) consistent with a role for SoxB1 factors in poising them for later expression (Sup. Fig. 9). Of the ∼33,000 Sox3 peaks associated with annotated genes, only 36% of these genes are expressed at blastula stage (Fig. 5E). GO term analysis of the 64% of genes not expressed in blastula stem cells showed enrichment for GO terms associated with central and peripheral nervous system development (Fig. 5F), cell types that SoxB1 factors play key roles in later in development (Stevanovic et al., 2021). Together these analyses provide evidence that SoxB1 factors may poise genes involved in the development of the neural plate border and other ectodermal lineages as early as blastula stages.

As our results demonstrated that Sox3 directly binds neural plate border genes in blastula stem cells, we next asked if SoxB1 factors regulate expression of these genes at this stage. We performed RNA-seq on dissected blastula stem cells (stage 9) depleted for *sox2* and *sox3*. DESeq2 was used to identify genes differentially expressed between *soxb1* morphant and control blastula stem cells. We found 2,235 genes to be differentially expressed (padj < 0.05) with approximately equal numbers up- and downregulated (Fig. 5G,H). Approximately 85% of differentially expressed genes had an associated blastula stage Sox3 ChIP peak (Fig. 5H). Among the genes downregulated in *soxb1* morphant cells were several canonical neural plate border factors, including *zic1*, *msx1*, and *tfap2a* (Fig. 5G). We also observed an up-regulation of *sox2* and *sox3* gene expression in *soxb1* morphants cells. These data suggest that not only do SoxB1 factors bind to regulatory regions of neural plate border genes, but they also promote expression of a subset of these genes at blastula stages.

### Pou5f3 factors are co-expressed with SoxB1 factors in neural plate border cells

SoxB1 factors require a transcriptional partner to promote gene expression (Kondoh and Kamachi, 2010). To identify candidate SoxB1 transcriptional partners involved in initiating and maintaining neural plate border gene expression, we identified Sox3 peaks that were shared between our Sox3 blastula stage and neural plate border (stage 11.5) ChIP-seq datasets. We found ∼3,500 shared peaks, representing 66% of the total peaks in the neural plate border dataset (Fig. 6A). Motif analysis of these shared peaks identified high enrichment for Sox factor, non-Sox HMG box factor (Lef/Tcf), and POU factor motifs. As we noted that an Oct4-Sox2 consensus sequence was present in approximately 25% of shared peaks (Fig. 6B; Sup. Fig. 10A), we examined if the Sox3 peaks associated with common neural plate border genes possessed Oct4-Sox2 consensus sequences. We found that most of these genes had at least one Oct4-Sox2 consensus site (Fig. 6C). We also performed motif analysis on shared regions of Sox3 enrichment for the genes within the four k-means clusters and found that POU motifs, including motifs for Oct4-Sox2, were enriched in the neural plate border cluster (Sup. Fig. 10B). These findings are consistent with recent ATAC-seq data which found enrichment for Oct4-Sox2 motifs in avian neural plate border/early neural crest cells (Hovland et al., 2022).

**Figure 6.**
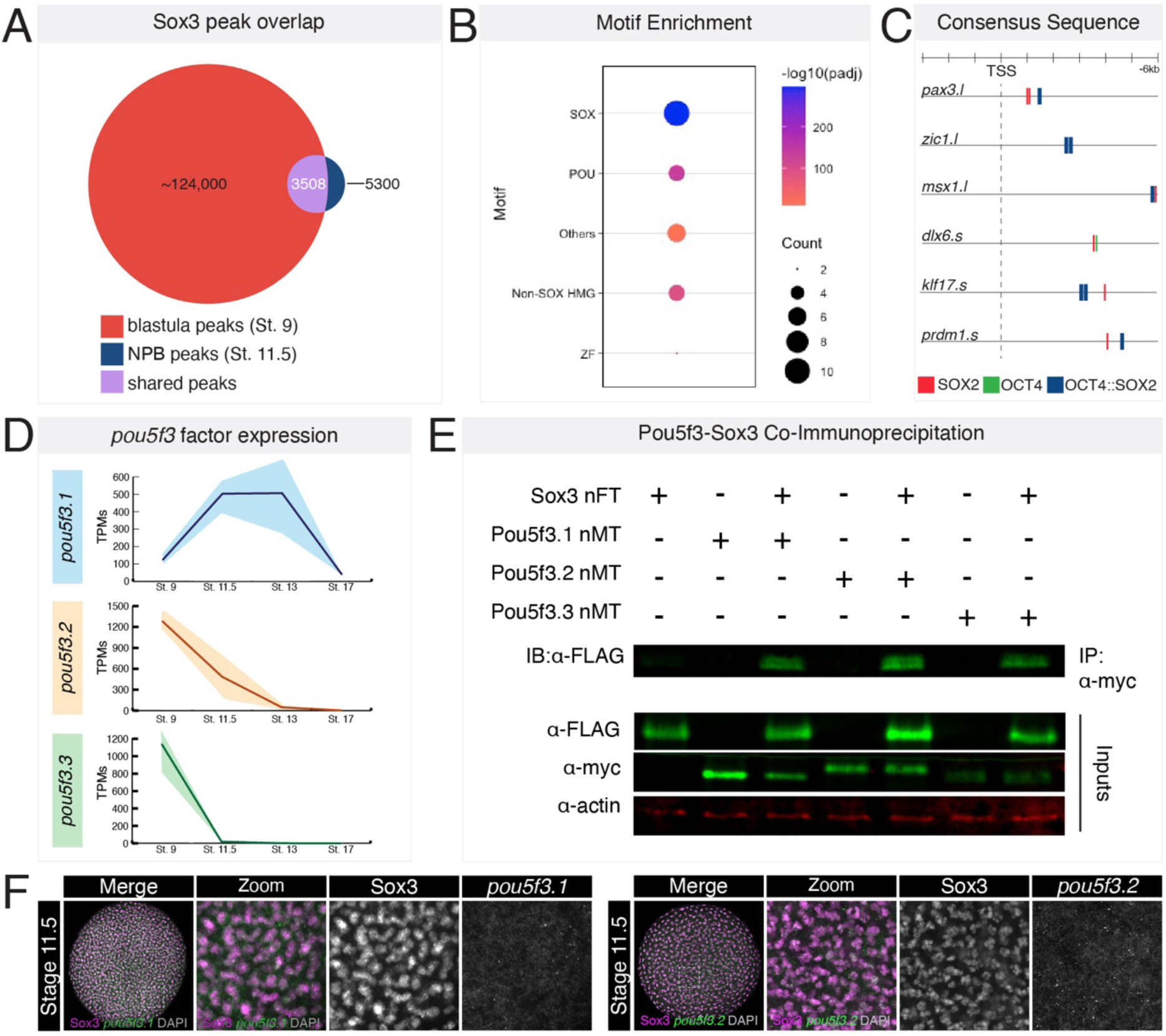
Motif analysis identifies POU factors as potential transcriptional partner involved in neural plate border formation. (A) Venn diagram showing overlap (purple) between Sox3 blastula stage peaks (red) and Sox3 neural plate border cell peaks (red). (B) Dot plot showing motif enrichment for shared category (purple) of Sox3 peaks. Motifs for POU transcription factors are highly enriched. (C) Localization of Sox2, Oct4, and Oct4::Sox2 motifs upstream of neural plate border genes transcription start sites. (D) TPM plots for *pou5f3* factor expression from blastula stem cells (stage 9), neural plate border-induced explants (stage 11.5, stage 13) and neural crest-induced explants (stage 17). (E) Co-immunoprecipitation of Sox3 with Pou5f3.1, Pou5f3.2, Pou5f3.3 factors. (F) Neural plate border-induced explants (stage 11.5) immunostained for Sox3 (magenta) and probed with HCR oligos for *pou5f3.1* or *pou5f3.2* (green). DAPI is shown in gray. n-terminal myc tag (nMT); n-terminal flag tag (nFT); zinc finger (ZF)

Given that combined activity of Pou5f (Oct3/4) and SoxB1 factors is important for maintenance of pluripotency in mouse and human embryonic stem cells (Dailey and Basilico, 2001), we wished to determine if they jointly regulate neural plate border formation. *Xenopus laevis* has three Pou5f3 homologs, *pou5f3.1*, *pou5f3.2*, and *pou5f3.3*.

*Pou5f3.3* is highly expressed in blastula stem cells, with low levels of expression in neural plate border cells, while *pou5f3.1* and *pou5f3.2* are expressed in both blastula stem cells and neural plate border cells (Fig. 6D) (York et al., in revision; Morrison and Brickman, 2006). Using co-immunoprecipitation experiments we found that all three Pou5f3 factors can physically interact with Sox3 (Fig. 6E). We next assessed if *pou5f3.1/2* and Sox3 are co-expressed in forming neural plate border cells. We used HCR probes for *pou5f3.1*/*pou5f3.2* and immunostaining for Sox3 and found co-expression of *pou5f3* and Sox3 in neural plate border-induced explants (Fig. 6F). These data indicate that SoxB1 factors and Pou5f3 factors are co-expressed in early neural plate border cells and could be functioning to regulate their formation.

### Combined activity of Sox3 and Pou5f3 is required for neural plate border formation

To determine if the combined activity of SoxB1 and Pou5f3 is required for neural plate border formation, we knocked down expression of *sox3*, *pou5f3.1*, and *pou5f3.2* using morpholinos and examined the expression of the neural plate border genes at the end of gastrulation. Both *pax3* and *zic1* expression was lost in triple morphant embryos (Fig. 7A), whereas loss of individual loss of *pou5f3.1/2* expands the neural plate border (York et al., in revision). Similarly, when we pharmacologically induced animal pole explants to a neural plate border state, we found that triple morphant explants were poorly/weakly induced compared to controls (Fig. 7B; Sup. Fig. 11), suggesting that the combined activity of SoxB1 and Pou5f3 transcription factors is essential for maintenance of the neural plate border gene expression.

**Figure 7.**
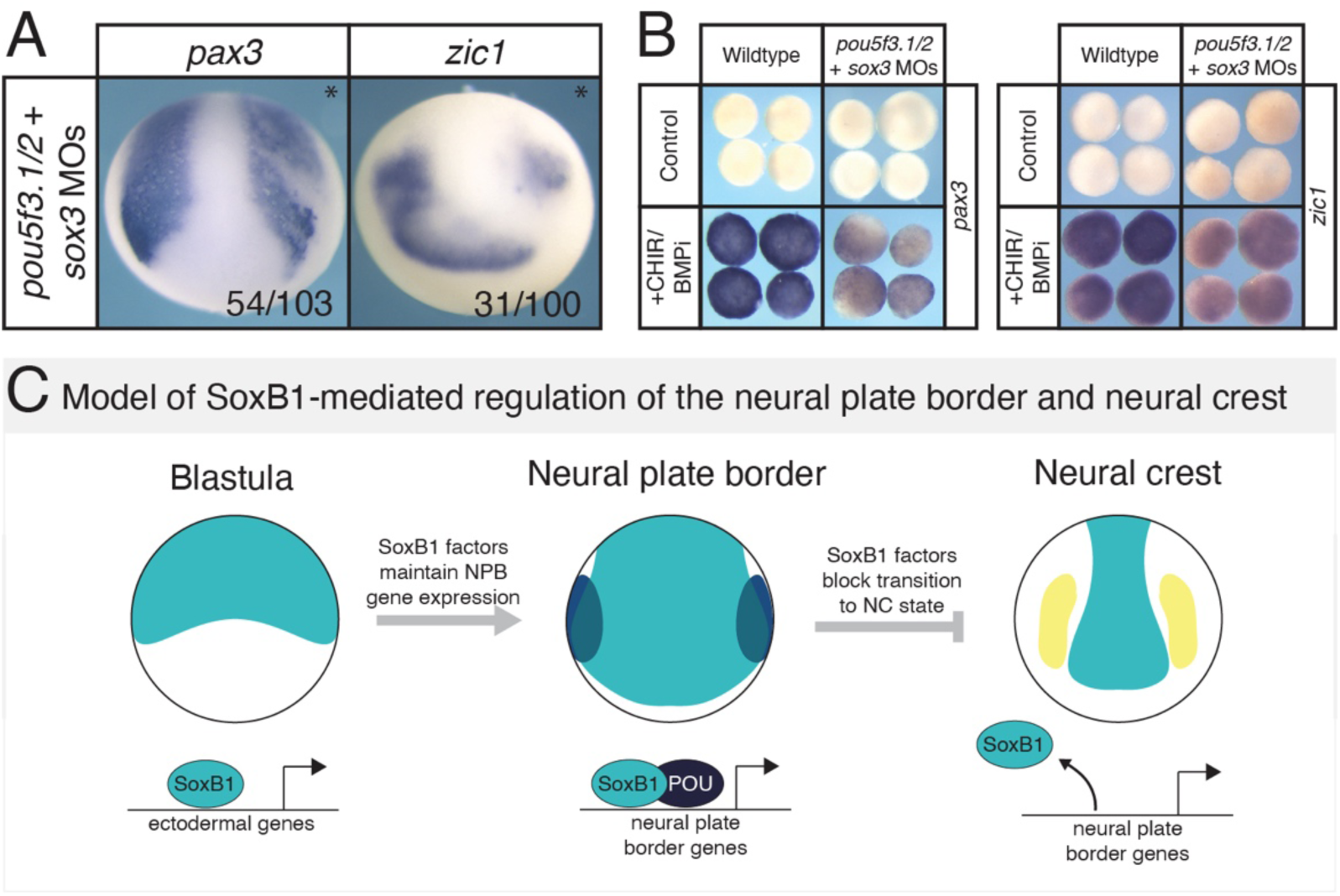
Combined activity of Pou5f3 and SoxB1 transcription factors is required for neural plate border formation. (A) *In situ* hybridization for pax3 and *zic1* at early neurula embryos injected unilaterally (injected side marked with *) with *sox3* + *pou5f3.1*+ *pou5f3.2* morpholinos. Fluorescein dextran was used as a lineage tracer and embryos were presorted for left/right side targeting. (B) *In situ* hybridization for *pax3* and *zic1* on wildtype and triple-morphant neural plate border-induced explants (stage 13). (C) Proposed model for SoxB1-mediated regulation of the neural plate border and neural crest. SoxB1 factors (teal) bind near ectodermal genes in blastula stem cells and promote or poise gene expression. SoxB1 factors, in partnership with Pou5f transcription factors (navy) maintain neural plate border gene (blue) expression throughout gastrulation. Downregulation of SoxB1 in neural plate border cells is required for neural crest (yellow) gene expression. Morpholino (MO); neural plate border (NPB); neural crest (NC)

## Discussion

We set out to determine if SoxB1 transcription factors play a role in establishing the neural plate border. Our ChIP-seq data provides evidence that SoxB1 factors may function to poise a broad range of ectodermal genes in blastula stem cells, including neural plate border genes. Through knockdown experiments, we determined that SoxB1 factors regulate the expression of neural plate border genes at blastula stages and are required for neural plate border formation. We also show that SoxB1 factors promote neural plate border gene expression and that downregulation of SoxB1 factors following neural plate border establishment is necessary for the transition from a neural plate border to a neural crest state. Our ChIP-seq data provide evidence that regulation of neural plate border gene expression by SoxB1 factors is likely direct. Finally, we demonstrate that combined activity of SoxB1 and Pou5f3 transcription factors is necessary for maintaining neural plate border gene expression (Fig. 7C).

### Transitioning from a neural plate border state to a neural crest state

Embryonic development relies upon precise regulation of developmental transitions to establish new cell types. Here we identify SoxB1 factors as negative regulators of the transition from a neural plate border to neural crest state. Prior gain- and loss-of-function experiments suggest there is a delicate balance between levels of neural plate border transcription factors and neural crest gene expression. In *Xenopus*, gain-of-function experiments for *pax3* and *zic1* have shown that while low levels of expression promote neural crest formation, at high levels they inhibit neural crest formation and promote hatching gland formation (Hong and Saint-Jeannet, 2007). We show here that expression of neural plate border genes is expanded and prolonged while neural crest formation is delayed in *soxb1* injected embryos. These results are similar to *pax3*/*zic1* gain-of-function experiments (high levels) and further emphasize the importance of strict temporal regulation of these neural plate border factors as well as maintenance of specific protein levels. It is also likely that neural plate border factors regulate the domain of *soxb1* expression, as *sox2* expression in the neural plate has been found to expand in *pax3* or *zic1* morphant embryos (Hong and Saint-Jeannet, 2007). Taken together, these findings support a model where SoxB1 factors promote low levels of neural plate border gene expression in blastula stem cells, and as those protein levels increase during neural plate border formation, they begin to restrict *soxb1* expression medially to the neural plate.

### Parallels between SoxB1-dependent transcriptional regulation in neural plate border and neural cell types

A major function of SoxB1 factors during development is to maintain a progenitor state and block differentiation. SoxB1 factors function in this manner in neural progenitor cells, where downregulation of SoxB1 factors promotes neuronal differentiation (Zhang and Cui, 2014). Here, we find another example where SoxB1 factors function to maintain a progenitor cell state – the neural plate border. Interestingly, several neural plate border genes are also expressed in neural cell types later in development (Inoue et al., 2007; Lin et al., 2016; Liu et al., 2004; Yu et al., 2011). In chick spinal cords, Zic1 and Msx1 block neuronal differentiation, promoting a progenitor state (Aruga et al., 2002b; Liu et al., 2004). Sox2 is also expressed in spinal cord neurons and functions to maintain pan-neural properties of neural progenitor cells (Graham et al., 2003). It is possible that SoxB1 factors may also function to directly regulate expression of these genes in neural cell types and together function to maintain a progenitor state.

### SoxB1-Pou5f3 regulation of neural plate border and neural crest formation

Sox and POU transcription factors directly interact to regulate many aspects of embryonic development. In neural progenitor cells, Sox2 and the class III POU factor, Brn2, synergize to activate *nestin* expression (Tanaka et al., 2004). Additionally, Sox2 and Oct4 (Pou V factor) are well known for regulating pluripotency in embryonic stem cells (Avilion et al., 2003; Boyer et al., 2005; Lodato et al., 2013). We identify another role for SoxB1 and Pou5f3 factors in regulation of neural plate border gene expression. Recent work in chick has identified a role for the SOX2-OCT4 heterodimer in establishing a neural crest epigenomic signature and showed that prolonged heterodimer expression of leads to maintenance of *PAX7* expression in what appears to be migrating neural crest cells (Hovland et al., 2022). While these data may seem to conflict with our findings, it is possible that prolonged *PAX7* expression is reflective of maintaining a neural plate border state rather than an early neural crest state. *PAX7* is one of the earliest markers for this cell population in chick (Prasad et al., 2020). Analysis of additional markers is necessary to distinguish between these two possibilities. Additionally, work in human cell culture has shown a global reduction in Sox2 binding in neural plate border cells relative to blastula cells (Hovland et al., 2022), which we also observe in *Xenopus*. However, we observed that Sox2 binds to neural plate border and neural crest genes in blastula stem cells whereas they observed SOX2-OCT4 binding to neural crest *cis*-regulatory regions upon neural crest specification *in vitro* (Hovland et al., 2022). ATAC-seq in mouse has shown that the *cis*-regulatory landscape of Oct4+ neural crest progenitor cells and epiblast stem cells are highly correlated, similar to our findings (Zalc et al., 2021). It is possible that noted differences in chromatin accessibility across these datasets is due to species-specific differences or a difference in timing of genome occupancy experiments. It is also worth noting that SoxB1 factors and Pou5f3 factors have different effects on neural crest gene expression. The neural crest is expanded in *pou5f3* injected *Xenopus* embryos (York et al., in revision), suggesting that Pou5f3 factors may have functions outside a SoxB1-Pou5f3 dimer during neural plate border/neural crest development. An interesting hypothesis is that Pou5f3 factors may be interacting with SoxE proteins in neural crest cells to promote neural crest gene expression. It is known that SoxB1 transcription factors interact with Pou5f3 factors through the highly conserved HMG domain (Remenyi et al., 2003), making it possible that Pou5f3 factors could also directly interact with SoxE factors, an area of current investigation.

### SoxB1 factors and the evolution of the neural crest

SoxB genes are present in the genome of basal metazoans and are hypothesized to have arisen in the last common ancestor of metazoans (Larroux et al., 2006). Both invertebrate chordates and vertebrates possess SoxB1 transcription factors (Heenan et al., 2016). The invertebrate chordate, amphioxus, possess lateral neural border cells, but lack neural crest. These cells are hypothesized to be the homolog to the vertebrate neural plate border and give rise to sensory neurons. In amphioxus embryos, *SoxB1* factors are expressed in the neural plate where expression persists in the neural tube until larval stages (Meulemans and Bronner-Fraser, 2007). Interestingly, SoxB1-a expression overlaps with lateral neural border gene expression in this species. Likewise, in lamprey, a jawless vertebrate, SoxB1a expression overlaps with neural plate border before restricting to the neural plate (Uy et al., 2012)(York et al., in revision). SoxB1 expressing cells are notably absent from the neural crest in lamprey, as in *Xenopus*, and are restricted to the neural tube (Uy et al., 2012). These expression data indicate that SoxB1 factors are expressed in the neural plate border in both invertebrate chordates and vertebrates, suggesting SoxB1-dependent regulation of neural plate border gene expression may be evolutionarily conserved. As amphioxus lack neural crest, it would be interesting to know SoxB1 if expression persists in lateral neural border cells. This could suggest that changes in SoxB1 *cis*-regulatory elements were needed for the downregulation of *SoxB1* expression in these cells for the neural crest to form. It is possible that modulation of SoxB1 expression in those cells may be one important distinguishing feature of the vertebrate vs nonvertebrate chordate border.

## Materials and Methods

### Embryological Methods

Wildtype *Xenopus laevis* embryos were obtained using standard methods and cultured in 0.1x Marc’s Modified Ringer’s Solution (MMR) (0.1 M NaCl, 2 mM KCl, 1 mM MgSO_4_, 2 mM CaCl_2_, 5 mM HEPES (pH 7.8), 0.1 mM EDTA) until desired stages (Nieuwkoop and Faber, 1994). Embryos or animal pole cell explants used for *in situ* hybridization, hybridization chain reaction (HCR), or immunofluorescence were fixed in 1x MEM (100mM MOPS (pH 7.4), 2mM EDTA, 1mM MgSO_4_) with 3.7% formaldehyde and dehydrated in methanol prior to use. Embryos or animal pole cell explants that underwent *in situ* hybridization were processed as described in (LaBonne and Bronner-Fraser, 1998) and imaged using an Infinity 8-8 camera (Teledyne Lumenera). Results are representative of a minimum of three biological replicates.

Microinjection of mRNA or morpholinos occurred at the 2-8 cell stage. mRNA was synthesized using a mMessage mMachine SP6 Transcription Kit (Invitrogen) and translation efficiency assessed by western blot. Either beta-galactosidase mRNA or fluorescein dextran were co-injected as a lineage tracer. Approximately 10-25 ng of translation-blocking morpholinos (Gene Tools) were injected per cell. Morpholino sequences are as follows:

*sox2*: 5’-TCTCCATCATGCTGTACAT-3’; *sox3*: 5’-TCCAACATGCTATACATTTGGAG-3’;

*pou5f3.1.L*: 5’-CCTATACAGCTCTTGCTCAAATC-3’;

*pou5f3.1.S*: 5’-GATTAAACATGATCTGTTGTCCG-3’;

*pou5f3.2.L*: 5’-CCAAGAGCTTGCAGTCAGATC-3’;

*pou5f3.2.S*: 5’-GCTGAACCCTAGAATGACCAG-3’

### Blastula explant assays

Animal pole cells were manually dissected using forceps at blastula stage (stage 9). Manipulated embryos were injected into either both cells at the 2-cell stage or the animal cells at the 4-8 cell stage with either mRNA or morpholino. To induce a neural plate border/neural crest state, dissected explants were immediately cultured in 3 μM K02288 (Sigma) and 107 μM CHIR99021 (Sigma) in 1xMMR, as described in (Huber and LaBonne, in revision) and remained in pharmacological solution until time of collection. To induce neural, dissected explants were immediately cultured in 20 μM K02288 (Sigma) and remained in pharmacological solution until time of collection.

### Western blot

Five whole embryos or ten explants were lysed in 1% NP-40 supplemented with protease inhibitors (Complete Mini, EDTA-free tablet (Roche), Leupeptin (Roche), Aprotinin (Sigma), and phenylmethylsulfonyl fluoride (PMSF; Sigma)). SDS page and western blot were used to detect proteins. The following primary antibodies were used: c-Myc 9E10 (1:3000; Santa Cruz; sc-40); FLAG M2 (1:3000; Sigma; F1804); actin (1:5000; Sigma; A2066); Sox3 (1:200, gift from Dominique Alfandari). IRDyes (1:20,000 mouse-800 CW; rabbit-680 TL) and the Odyssey platform (LI-COR Biosciences) were used to detect proteins. Image Studio Lite software was used to quantify protein. Results are representative of a minimum of three biological replicates.

#### Co-immunoprecipitation

Five whole embryos were lysed in 1% NP-40 supplemented with protease inhibitors (see above). A five percent input was retained for western blot analysis and the remaining 95% incubated with c-Myc 9E10 antibody (1:500) for one hour. Approximately 25-30 μl of PAS beads (Sigma; P3391) were added to lysate and incubated for two hours. Beads were washed with 1% NP-40 and remaining proteins eluted off the beads. Input and immunoprecipitation samples were analyzed via western blot as described above. Results are representative of a minimum of three biological replicates.

### RNA isolation, library prep, and sequencing

RNA was extracted from 10-12 explants using Trizol (Life Technologies) followed by LiCl precipitation. Approximately 300 ng of RNA was used for library prep (NEBNext® Ultra™ II Directional RNA Library Prep Kit for Illumina) following standard kit procedure. Libraries were pooled and sequence using NextSeq 500 system (Single end, 75 bp). Results are representative of three biological replicates.

### Quantitative PCR

RNA was isolated as described above and converted into cDNA using a High-Capacity cDNA Reverse Transcription Kit (Life Technologies). qPCR was performed using SYBR Premix (Clontech #RR820W) using the below primer sequences. Fold expression was normalized to ornithine decarboxylase (*odc*) and relative to wildtype. The ΔΔCT method was used to calculate fold expression and represented as a mean from three separate biological replicates with error bars representing standard deviation.

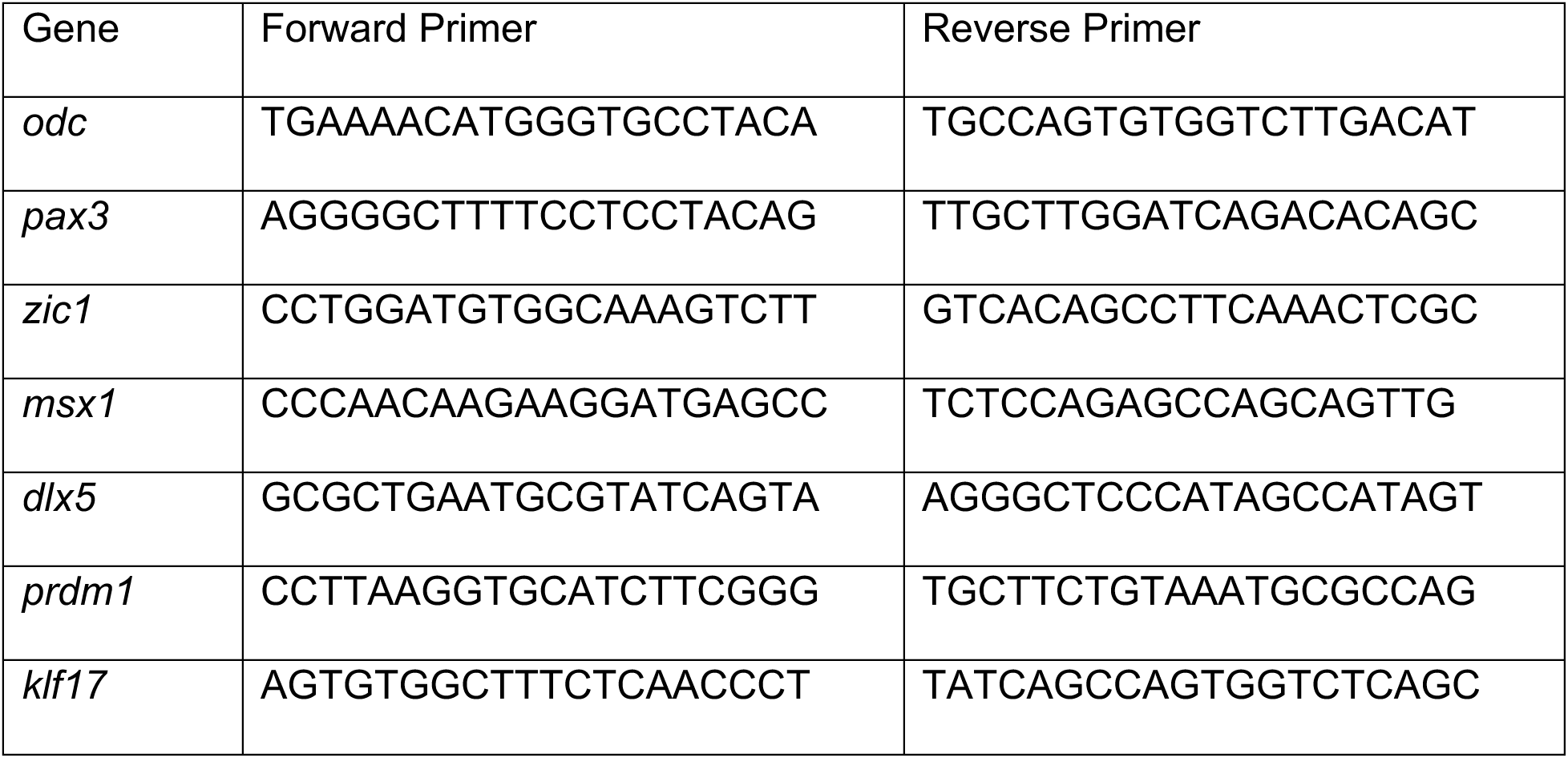

### Hybridization Chain Reaction and Immunofluorescence

Hybridization chain reaction (HCR) methodologies are slightly modified from (Choi et al., 2018). Whole embryos or explants were hybridized with DNA probe sets for *pax3* (exonic and intronic), *sox10*, foxd3, *pou5f3*.*1*, *pou5f3*.*2* (Molecular Instruments) and incubated overnight at 37 °C. Probe was removed, samples washed, and then incubated overnight with DNA hairpins labeled with Alexa 647 or Alexa 546 (Molecular Instruments). Unbound hairpins were removed via 5x SSC washes and then samples were immediately blocked in 10% fetal bovine serum with 0.1% triton (in PBS) in prep for Sox3 immunostaining. Samples were incubated for 1 hour at room temp with Sox3 antibody (1:100, gift from Dominique Alfandari) or c-Myc 9E10 (1:1000), washed, and incubated for 1 hour with Alexa 488/568 secondary (1:1000; Life Technologies, mouse) and DAPI (1:5000; Life Technologies). Sox2 antibody was used at 1: 500 (Cell Signaling Technologies; 3579S). Samples were mounted and imaged using a Nikon C2 upright confocal with two GaAsP detectors and four standard laser lines.

### Co-localization Analysis

Max intensity projections of explants were made from confocal files using Fiji. Sox3 and *pax3* expression was autothresholded using the RenyiEntropy method, and DAPI was auto local thresholded using the Bernsen method using Fiji. To discern real cells, only connected clusters of 20 DAPI-expressing pixels or larger were selected to be analyzed. DAPI-expressing cells were quantified and used as an overlay to quantify *pax3* and Sox3-positive cells in those respective channels. The number of *pax3*-postive cells, Sox3-positive cells, and p*ax3*/Sox3 double-positive cells were quantified. Three regions in each explant, with 50 DAPI-positive cells in each region, were randomly selected for further analysis. Within each region, the total number of *pax3-*postive cells that were also Sox3-positive (Sox3&*pax3*++/total *pax3*+ cells) were quantified. 18 regions in total were analyzed per stage (three replicates; two explants per stage; three regions per explant; total sample size: 18). All values were visualized via violin plots (ggplot2) using the Tukey method to define outliers.

### Chromatin Immunoprecipitation (ChIP)

Fifty wildtype or myc-tagged Sox3 expressing animal pole cell explants were crosslinked for 15 minutes with 1% methanol-free formaldehyde (Life Technologies) and the reaction was quenched with 125 mM glycine in 0.5% PBST (Triton X-100). Explants were washed with 0.01x MMR then lysed using ChIP lysis buffer (5mM Tris-HCl, pH 7.4, 15 mM NaCl, 1mM EDTA, 1mM DTT, 1% NP-40, 0.25% Sodium deoxycholate, 0.1% SDS with protease inhibitors). Lysates were briefly sonicated using a M220 focused ultrasonicator and incubated with Myc antibody (1:500; Sigma, C3956) overnight at 4 °C. Dynabeads Protein G (Life Technologies; 10004D) were added to samples and incubated for 1 hour. Beads were washed with Tagmentation Wash Buffer (10mM Tris, pH 7.5, 5mM MgCl_2_) and incubated with Tn5 transposes (Illumina Tagment DNA Enzyme and Buffer Small Kit) for 40 minutes at 37 °C in a thermomixer (1000 rpm). Tagmented samples were washed, eluted from beads, reverse crosslinked, and purified using Qiagen MinElute Reaction Clean Up kit (#28204). DNA was amplified using Nextera primers, adapter dimers removed with AMPure XP beads (Beckman), and library quality assessed via Tapestation. Pooled libraries were sequenced using NextSeq 500 (blastula) or NovaSeq (neural plate border) systems (Paired end, 75 bp). Results are representative of three biological replicates.

### DNA constructs

*Xenopus laevis sox2* and *sox3* clones were obtained from Open Biosystems and subcloned into a pCS2+ vector that add either five N-or C-terminal Myc tags (nMT/cMT). All constructs were verified via sequencing. *Xenopus laevis pou5f3.1/2/3* constructs are described in (York et al., in revision).

### RNA-seq analysis

Fastq files were obtained from Northwestern Sequencing core. Sequences were trimmed using fastp (Chen et al., 2018) and aligned to the *Xenopus laevis* genome (9.2) using STAR or RSEM (for TPMs) (Dobin et al., 2013; Li and Dewey, 2011). TPM data is an average of three biological replicates of combined data from S and L alleles for each gene, the width of each line represents standard deviation. HTseq was used to generate read counts from STAR aligned bam files (Putri et al., 2022). Differential expression analysis was performed using DESeq2 with significance defined as *p*_adj_ ≤ 0.05 (Love et al., 2014). Volcano plots were generated using ggplot2 (Wickham, 2016).

### ChIP-seq analysis

Fastq files were obtained from Northwestern Sequencing core. Sequences were trimmed using fastp (Chen et al., 2018) and aligned to the *Xenopus laevis* genome (9.2) using bowtie2 (Langmead and Salzberg, 2012). Samtools was used to process files (Danecek et al., 2021). Name sorted bam files were used for peak calling using genrich (-t Sox3 bam files; -c wildtype bam files; -v; -e MT; -a 20) (available at https://github.com/jsh58/Genrich). HOMER was used to annotate peaks and for motif analysis (Heinz et al., 2010). The FIMO tool from MEME Suite was also used for motif analysis (Grant et al., 2011). The GO Consortium was used for GO term analysis, using the *Homo sapiens* database (Ashburner et al., 2000; Gene Ontology, 2021). IGV was used for browser track visualization (Robinson et al., 2011).

### Statistics

Fisher’s exact test was performed on 2×2 contingency plots and odds ratios visualized by forest plots generated in ggplot2. All other statistical analyses used two-tailed t-tests to determine significance.

### Animals

All animal procedures were approved by the Institutional Animal Care and Use Committee, Northwestern University, and are in accordance with the National Institutes of Health’s Guide for the Care and Use of Laboratory Animals.

## Acknowledgements

The authors would like to thank Shelby Blythe (Northwestern University) for valuable insights on ChIP-seq procedures/analyses and for critical reading of the manuscript. We would also like to thank Paul Huber for sharing pharmacological induction protocols, Andrew Montequin for HCR protocols and members of the LaBonne lab for helpful discussions.

## References

Ariizumi, T., Asashima, M., 2001. In vitro induction systems for analyses of amphibian organogenesis and body patterning. The International journal of developmental biology 45, 273–279.

Aruga, J., Inoue, T., Hoshino, J., Mikoshiba, K., 2002a. Zic2 controls cerebellar development in cooperation with Zic1. The Journal of neuroscience : the official journal of the Society for Neuroscience 22, 218–225.

Aruga, J., Tohmonda, T., Homma, S., Mikoshiba, K., 2002b. Zic1 promotes the expansion of dorsal neural progenitors in spinal cord by inhibiting neuronal differentiation. Developmental biology 244, 329–341.

Ashburner, M., Ball, C.A., Blake, J.A., Botstein, D., Butler, H., Cherry, J.M., Davis, A.P., Dolinski, K., Dwight, S.S., Eppig, J.T., Harris, M.A., Hill, D.P., Issel-Tarver, L., Kasarskis, A., Lewis, S., Matese, J.C., Richardson, J.E., Ringwald, M., Rubin, G.M., Sherlock, G., 2000. Gene ontology: tool for the unification of biology. The Gene Ontology Consortium. Nature genetics 25, 25–29.

Avilion, A.A., Nicolis, S.K., Pevny, L.H., Perez, L., Vivian, N., Lovell-Badge, R., 2003. Multipotent cell lineages in early mouse development depend on SOX2 function. Genes & development 17, 126–140.

Boyer, L.A., Lee, T.I., Cole, M.F., Johnstone, S.E., Levine, S.S., Zucker, J.P., Guenther, M.G., Kumar, R.M., Murray, H.L., Jenner, R.G., Gifford, D.K., Melton, D.A., Jaenisch, R., Young, R.A., 2005. Core transcriptional regulatory circuitry in human embryonic stem cells. Cell 122, 947–956.

Buitrago-Delgado, E., Nordin, K., Rao, A., Geary, L., LaBonne, C., 2015. NEURODEVELOPMENT. Shared regulatory programs suggest retention of blastula-stage potential in neural crest cells. Science 348, 1332–1335.

Buitrago-Delgado, E., Schock, E.N., Nordin, K., LaBonne, C., 2018. A transition from SoxB1 to SoxE transcription factors is essential for progression from pluripotent blastula cells to neural crest cells. Developmental biology.

Bylund, M., Andersson, E., Novitch, B.G., Muhr, J., 2003. Vertebrate neurogenesis is counteracted by Sox1-3 activity. Nat Neurosci 6, 1162–1168.

Cattell, M.V., Garnett, A.T., Klymkowsky, M.W., Medeiros, D.M., 2012. A maternally established SoxB1/SoxF axis is a conserved feature of chordate germ layer patterning. Evol Dev 14, 104–115.

Chen, S., Zhou, Y., Chen, Y., Gu, J., 2018. fastp: an ultra-fast all-in-one FASTQ preprocessor. Bioinformatics 34, i884–i890.

Choi, H.M.T., Schwarzkopf, M., Fornace, M.E., Acharya, A., Artavanis, G., Stegmaier, J., Cunha, A., Pierce, N.A., 2018. Third-generation in situ hybridization chain reaction: multiplexed, quantitative, sensitive, versatile, robust. Development 145.

Dailey, L., Basilico, C., 2001. Coevolution of HMG domains and homeodomains and the generation of transcriptional regulation by Sox/POU complexes. J Cell Physiol 186, 315–328.

Danecek, P., Bonfield, J.K., Liddle, J., Marshall, J., Ohan, V., Pollard, M.O., Whitwham, A., Keane, T., McCarthy, S.A., Davies, R.M., Li, H., 2021. Twelve years of SAMtools and BCFtools. Gigascience 10.

Dobin, A., Davis, C.A., Schlesinger, F., Drenkow, J., Zaleski, C., Jha, S., Batut, P., Chaisson, M., Gingeras, T.R., 2013. STAR: ultrafast universal RNA-seq aligner. Bioinformatics 29, 15–21.

Garnett, A.T., Square, T.A., Medeiros, D.M., 2012. BMP, Wnt and FGF signals are integrated through evolutionarily conserved enhancers to achieve robust expression of Pax3 and Zic genes at the zebrafish neural plate border. Development 139, 4220–4231.

Gene Ontology, C., 2021. The Gene Ontology resource: enriching a GOld mine. Nucleic acids research 49, D325–D334.

Godichon-Baggioni, A., Maugis-Rabusseau, C., Rau, A., 2019. Clustering transformed compositional data using K-means, with applications in gene expression and bicycle sharing system data. Journal of Applied Statistics 46, 47–65.

Graham, V., Khudyakov, J., Ellis, P., Pevny, L., 2003. SOX2 functions to maintain neural progenitor identity. Neuron 39, 749–765.

Grant, C.E., Bailey, T.L., Noble, W.S., 2011. FIMO: scanning for occurrences of a given motif. Bioinformatics 27, 1017–1018.

Groves, A.K., LaBonne, C., 2014. Setting appropriate boundaries: fate, patterning and competence at the neural plate border. Developmental biology 389, 2–12.

Heenan, P., Zondag, L., Wilson, M.J., 2016. Evolution of the Sox gene family within the chordate phylum. Gene 575, 385–392.

Heinz, S., Benner, C., Spann, N., Bertolino, E., Lin, Y.C., Laslo, P., Cheng, J.X., Murre, C., Singh, H., Glass, C.K., 2010. Simple combinations of lineage-determining transcription factors prime cis-regulatory elements required for macrophage and B cell identities. Mol Cell 38, 576–589.

Hong, C.S., Saint-Jeannet, J.P., 2007. The activity of Pax3 and Zic1 regulates three distinct cell fates at the neural plate border. Molecular biology of the cell 18, 2192–2202.

Horr, B., Kurtz, R., Pandey, A., Hoffstrom, B.G., Schock, E., LaBonne, C., Alfandari, D., 2023. Production and characterization of monoclonal antibodies to Xenopus proteins. Development 150.

Hovland, A.S., Bhattacharya, D., Azambuja, A.P., Pramio, D., Copeland, J., Rothstein, M., Simoes-Costa, M., 2022. Pluripotency factors are repurposed to shape the epigenomic landscape of neural crest cells. Developmental cell 57, 2257–2272 e2255.

Inoue, T., Ota, M., Ogawa, M., Mikoshiba, K., Aruga, J., 2007. Zic1 and Zic3 regulate medial forebrain development through expansion of neuronal progenitors. The Journal of neuroscience : the official journal of the Society for Neuroscience 27, 5461–5473.

Johnson, K., Freedman, S., Braun, R., LaBonne, C., 2022. Quantitative analysis of transcriptome dynamics provides novel insights into developmental state transitions. BMC Genomics 23, 723.

Kishi, M., Mizuseki, K., Sasai, N., Yamazaki, H., Shiota, K., Nakanishi, S., Sasai, Y., 2000. Requirement of Sox2-mediated signaling for differentiation of early Xenopus neuroectoderm. Development 127, 791–800.

Kondoh, H., Kamachi, Y., 2010. SOX-partner code for cell specification: Regulatory target selection and underlying molecular mechanisms. The international journal of biochemistry & cell biology 42, 391–399.

LaBonne, C., Bronner-Fraser, M., 1998. Neural crest induction in Xenopus: evidence for a two-signal model. Development 125, 2403–2414.

Langmead, B., Salzberg, S.L., 2012. Fast gapped-read alignment with Bowtie 2. Nat Methods 9, 357–359.

Larroux, C., Fahey, B., Liubicich, D., Hinman, V.F., Gauthier, M., Gongora, M., Green, K., Worheide, G., Leys, S.P., Degnan, B.M., 2006. Developmental expression of transcription factor genes in a demosponge: insights into the origin of metazoan multicellularity. Evol Dev 8, 150–173.

Li, B., Dewey, C.N., 2011. RSEM: accurate transcript quantification from RNA-Seq data with or without a reference genome. BMC Bioinformatics 12, 323.

Lin, J., Wang, C., Yang, C., Fu, S., Redies, C., 2016. Pax3 and Pax7 interact reciprocally and regulate the expression of cadherin-7 through inducing neuron differentiation in the developing chicken spinal cord. J Comp Neurol 524, 940–962.

Liu, Y., Helms, A.W., Johnson, J.E., 2004. Distinct activities of Msx1 and Msx3 in dorsal neural tube development. Development 131, 1017–1028.

Lodato, M.A., Ng, C.W., Wamstad, J.A., Cheng, A.W., Thai, K.K., Fraenkel, E., Jaenisch, R., Boyer, L.A., 2013. SOX2 co-occupies distal enhancer elements with distinct POU factors in ESCs and NPCs to specify cell state. PLoS genetics 9, e1003288.

Love, M.I., Huber, W., Anders, S., 2014. Moderated estimation of fold change and dispersion for RNA-seq data with DESeq2. Genome Biol 15, 550.

Marchal, L., Luxardi, G., Thome, V., Kodjabachian, L., 2009. BMP inhibition initiates neural induction via FGF signaling and Zic genes. Proceedings of the National Academy of Sciences of the United States of America 106, 17437–17442.

Meulemans, D., Bronner-Fraser, M., 2007. The amphioxus SoxB family: implications for the evolution of vertebrate placodes. Int J Biol Sci 3, 356–364.

Monsoro-Burq, A.H., Wang, E., Harland, R., 2005. Msx1 and Pax3 cooperate to mediate FGF8 and WNT signals during Xenopus neural crest induction. Developmental cell 8, 167–178.

Morrison, G.M., Brickman, J.M., 2006. Conserved roles for Oct4 homologues in maintaining multipotency during early vertebrate development. Development 133, 2011–2022.

Nieuwkoop, P.D., Faber, J., 1994. Normal table of Xenopus laevis (Daudin) : a systematical and chronological survey of the development from the fertilized egg till the end of metamorphosis. Garland Pub., New York.

Okuda, Y., Yoda, H., Uchikawa, M., Furutani-Seiki, M., Takeda, H., Kondoh, H., Kamachi, Y., 2006. Comparative genomic and expression analysis of group B1 sox genes in zebrafish indicates their diversification during vertebrate evolution. Developmental dynamics : an official publication of the American Association of Anatomists 235, 811–825.

Papanayotou, C., Mey, A., Birot, A.M., Saka, Y., Boast, S., Smith, J.C., Samarut, J., Stern, C.D., 2008. A mechanism regulating the onset of Sox2 expression in the embryonic neural plate. PLoS biology 6, e2.

Pla, P., Monsoro-Burq, A.H., 2018. The neural border: Induction, specification and maturation of the territory that generates neural crest cells. Developmental biology 444 Suppl 1, S36–S46.

Prasad, M.S., Uribe-Querol, E., Marquez, J., Vadasz, S., Yardley, N., Shelar, P.B., Charney, R.M., Garcia-Castro, M.I., 2020. Blastula stage specification of avian neural crest. Developmental biology 458, 64–74.

Putri, G.H., Anders, S., Pyl, P.T., Pimanda, J.E., Zanini, F., 2022. Analysing high-throughput sequencing data in Python with HTSeq 2.0. Bioinformatics 38, 2943–2945.

Rau, A., Maugis-Rabusseau, C., 2018. Transformation and model choice for RNA-seq co-expression analysis. Brief Bioinform 19, 425–436.

Remenyi, A., Lins, K., Nissen, L.J., Reinbold, R., Scholer, H.R., Wilmanns, M., 2003. Crystal structure of a POU/HMG/DNA ternary complex suggests differential assembly of Oct4 and Sox2 on two enhancers. Genes & development 17, 2048–2059.

Rex, M., Orme, A., Uwanogho, D., Tointon, K., Wigmore, P.M., Sharpe, P.T., Scotting, P.J., 1997. Dynamic expression of chicken Sox2 and Sox3 genes in ectoderm induced to form neural tissue. Developmental dynamics : an official publication of the American Association of Anatomists 209, 323–332.

Robinson, J.T., Thorvaldsdottir, H., Winckler, W., Guttman, M., Lander, E.S., Getz, G., Mesirov, J.P., 2011. Integrative genomics viewer. Nat Biotechnol 29, 24–26.

Roellig, D., Tan-Cabugao, J., Esaian, S., Bronner, M.E., 2017. Dynamic transcriptional signature and cell fate analysis reveals plasticity of individual neural plate border cells. eLife 6.

Rogers, C.D., Harafuji, N., Archer, T., Cunningham, D.D., Casey, E.S., 2009. Xenopus Sox3 activates sox2 and geminin and indirectly represses Xvent2 expression to induce neural progenitor formation at the expense of non-neural ectodermal derivatives. Mechanisms of development 126, 42–55.

Sasai, Y., De Robertis, E.M., 1997. Ectodermal patterning in vertebrate embryos. Developmental biology 182, 5–20.

Sato, T., Sasai, N., Sasai, Y., 2005. Neural crest determination by co-activation of Pax3 and Zic1 genes in Xenopus ectoderm. Development 132, 2355–2363.

Silver, N., Cotroneo, E., Proctor, G., Osailan, S., Paterson, K.L., Carpenter, G.H., 2008. Selection of housekeeping genes for gene expression studies in the adult rat submandibular gland under normal, inflamed, atrophic and regenerative states. BMC Mol Biol 9, 64.

Soufi, A., Garcia, M.F., Jaroszewicz, A., Osman, N., Pellegrini, M., Zaret, K.S., 2015. Pioneer transcription factors target partial DNA motifs on nucleosomes to initiate reprogramming. Cell 161, 555–568.

Stevanovic, M., Drakulic, D., Lazic, A., Ninkovic, D.S., Schwirtlich, M., Mojsin, M., 2021. SOX Transcription Factors as Important Regulators of Neuronal and Glial Differentiation During Nervous System Development and Adult Neurogenesis. Front Mol Neurosci 14, 654031.

Tanaka, S., Kamachi, Y., Tanouchi, A., Hamada, H., Jing, N., Kondoh, H., 2004. Interplay of SOX and POU factors in regulation of the Nestin gene in neural primordial cells. Molecular and cellular biology 24, 8834–8846.

Theveneau, E., Mayor, R., 2012. Neural crest migration: interplay between chemorepellents, chemoattractants, contact inhibition, epithelial-mesenchymal transition, and collective cell migration. Wiley interdisciplinary reviews. Developmental biology 1, 435–445.

Tribulo, C., Aybar, M.J., Nguyen, V.H., Mullins, M.C., Mayor, R., 2003. Regulation of Msx genes by a Bmp gradient is essential for neural crest specification. Development 130, 6441–6452.

Uy, B.R., Simoes-Costa, M., Sauka-Spengler, T., Bronner, M.E., 2012. Expression of Sox family genes in early lamprey development. The International journal of developmental biology 56, 377–383.

Wakamatsu, Y., Endo, Y., Osumi, N., Weston, J.A., 2004. Multiple roles of Sox2, an HMG-box transcription factor in avian neural crest development. Developmental dynamics : an official publication of the American Association of Anatomists 229, 74–86.

Wickham, H., 2016. ggplot2 : Elegant Graphics for Data Analysis, Use R!,, 2nd ed. Springer International Publishing : Imprint: Springer,, Cham, pp. 1 online resource (XVI, 260 pages 232 illustrations, 140 illustrations in color.

Wilson, P.A., Lagna, G., Suzuki, A., Hemmati-Brivanlou, A., 1997. Concentration-dependent patterning of the Xenopus ectoderm by BMP4 and its signal transducer Smad1. Development 124, 3177–3184.

Yu, M., Xi, Y., Pollack, J., Debiais-Thibaud, M., Macdonald, R.B., Ekker, M., 2011. Activity of dlx5a/dlx6a regulatory elements during zebrafish GABAergic neuron development. Int J Dev Neurosci 29, 681–691.

Zalc, A., Sinha, R., Gulati, G.S., Wesche, D.J., Daszczuk, P., Swigut, T., Weissman, I.L., Wysocka, J., 2021. Reactivation of the pluripotency program precedes formation of the cranial neural crest. Science 371.

Zhang, S., Cui, W., 2014. Sox2, a key factor in the regulation of pluripotency and neural differentiation. World J Stem Cells 6, 305–311.

