## Supplemental Figures for "SoxB1 transcription factors are essential for initiating and maintaining the neural plate border gene expression"

Supplemental Figure 1

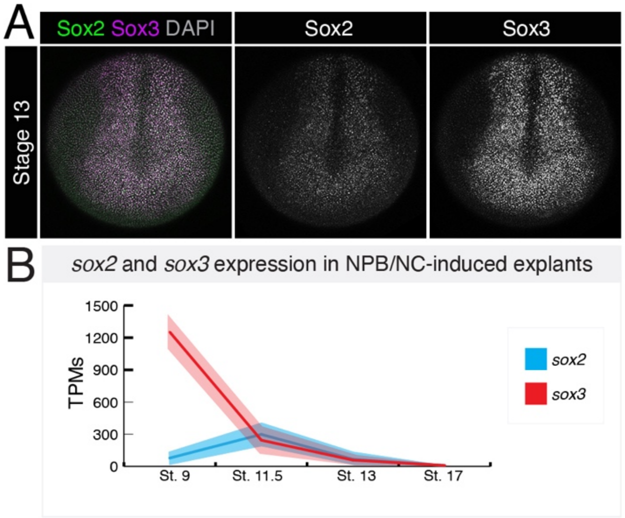


**Sup. Fig.1. Sox2 and Sox3 have overlapping expression domains.** (A) Wildtype stage 13 embryo immunostained for Sox2 (green) and Sox3 (magenta). DAPI is shown in gray. (B) TPMs for *sox2* (blue) and *sox3* (red) in in blastula stem cells (St. 9) and neural plate border/neural crest-induced explants (St. 11.5, St. 13, St. 17).

Supplemental Figure 2

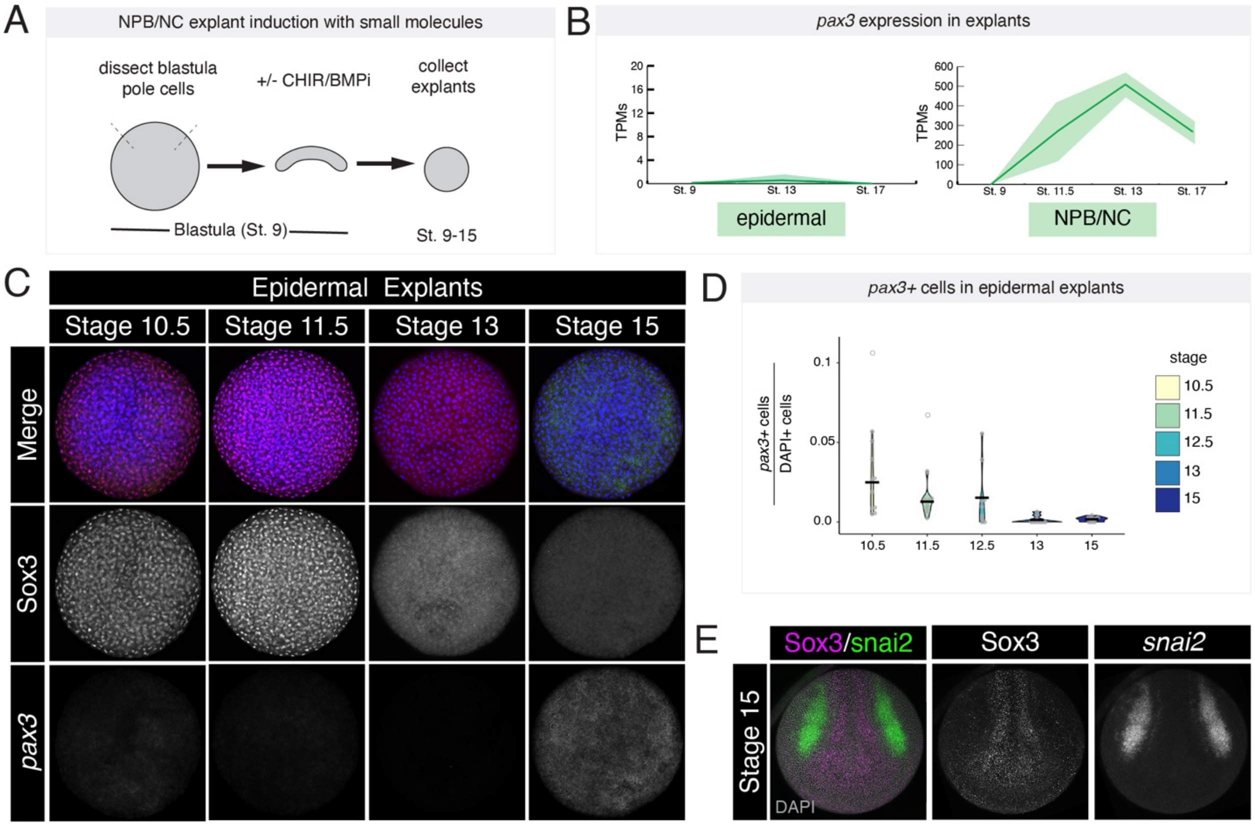


**Sup. Fig. 2. Temporal relationship between Sox3 and neural plate border gene expression.** (A) Experimental set up for neural plate border/neural crest induction of blastula pole cell explants using small molecules. (B) TPMs for pax3 in epidermal vs neural crest-induced explants (St. 9, 11.5, St. 13, St. 17). (C) Nascent *pax3* expression (green) and *Sox3* protein (red) in epidermal (control) explants from early gastrulation through mid-neurulation. DAPI is shown in blue. (D) Quantification of percent *pax3*+ cells in epidermal explants. (E) Wildtype stage 15 embryo immunostained for Sox3 (magenta) and probed for *snai2* (green) using HCR. DAPI is shown in gray.

Supplemental Figure 3

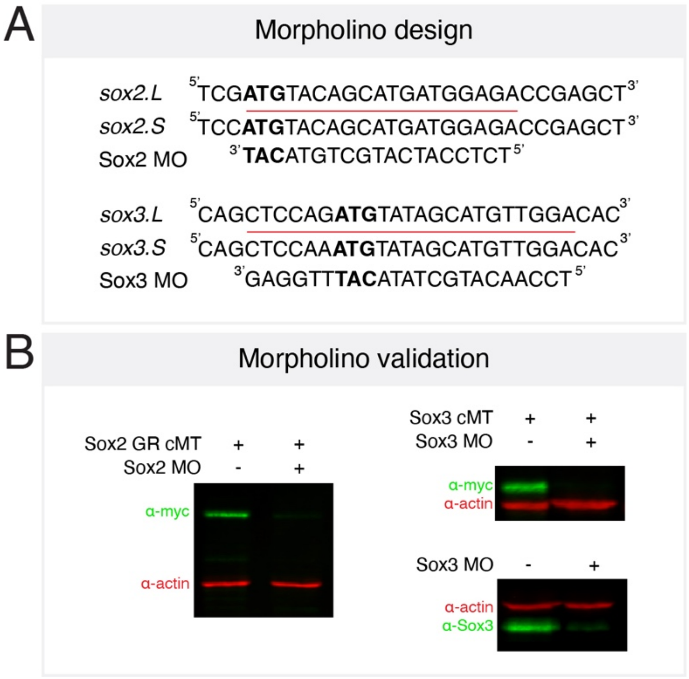


**Sup. Fig. 3. Sox2 and Sox3 morpholino validation.** (A) Schematic showing morpholino sequences and target regions at *sox2* and *sox3* alloalleles. (B) Western blot validation of sox2 and sox3 morpholinos. C-terminal myc tag (cMT); morpholino (MO)

Supplemental Figure 4

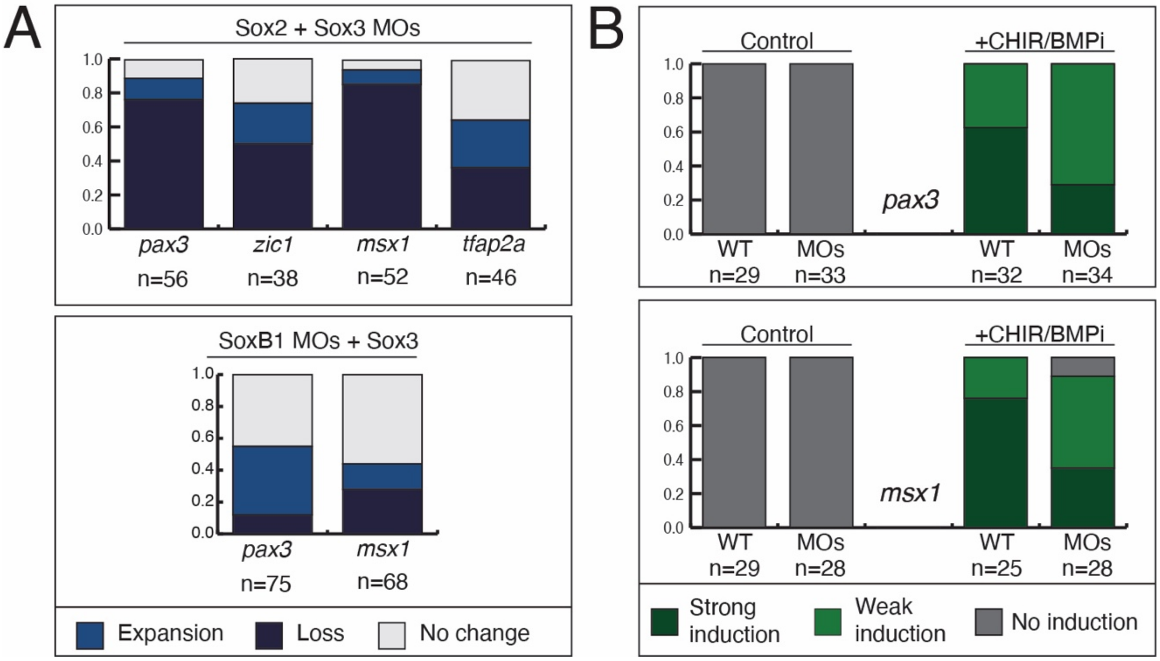


**Sup. Fig. 4. *soxb1* morphant scoring.** (A) Stacked bar graphs with the percent of embryos with changes in gene expression (loss, expansion, no change) for *sox2* and *sox3* double morphants and rescued morphants (B) Stacked bar graphs with the percent of neural plate border-induced explants expressing *pax3* or *msx1*, indicating induction to a neural plate border state. Morpholino (MO)

Supplemental Figure 5

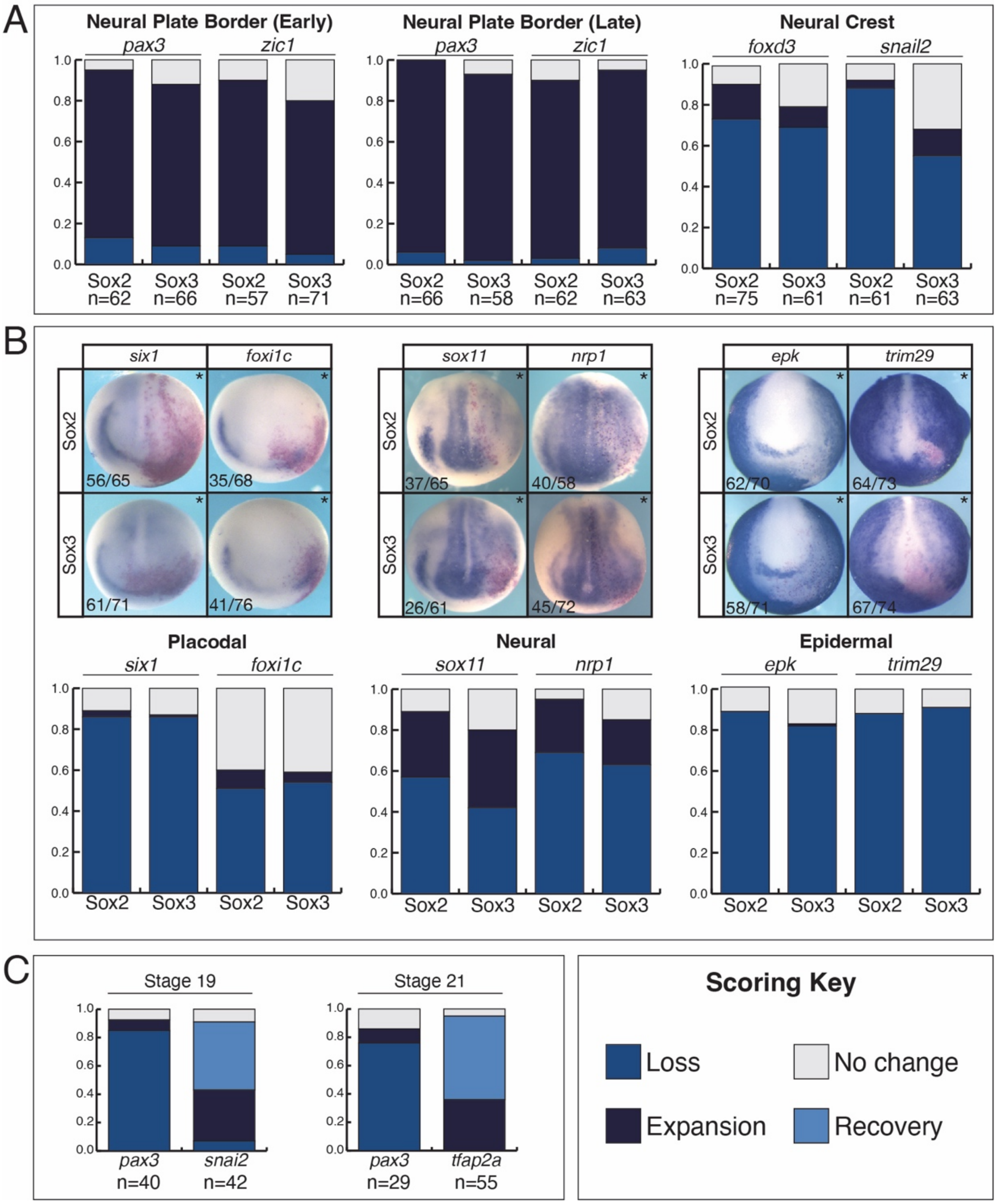


**Sup. Fig. 5. *In situ* hybridization and scoring for Sox2 and Sox3 expressing embryos.**

(A) Stacked bar graphs with the percent of embryos with changes in gene expression (loss, expansion, no change) in *sox2* or *sox3* expressing embryos. (B) *In situ* hybridization in stage 16 embryos unilaterally expressing *sox2* or *sox3* mRNA (* denotes injected side) probing for placodal markers (*six1* and *foxi1c),* neural markers (*sox11* and *nrp1)*, and epidermal markers (*epk* and *trim29*) with associated scoring. (D) Stacked bar graphs with the percent of sox3 expressing embryos with changes in gene expression (loss, expansion, recovery, no change) at stage 19 and 21.

Supplemental Figure 6

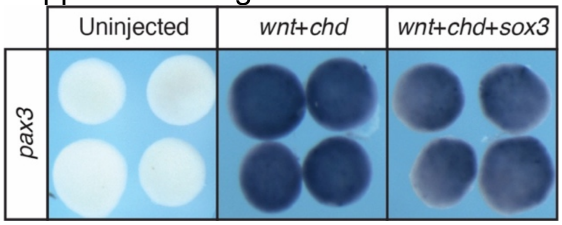


**Sup. Fig. 6. Neural plate border induction assessment for ChIP-seq experiments.** *In situ hybridization* for *pax3* in neural plate border-induced explants with and without myc-tagged sox3.

Supplemental Figure 7

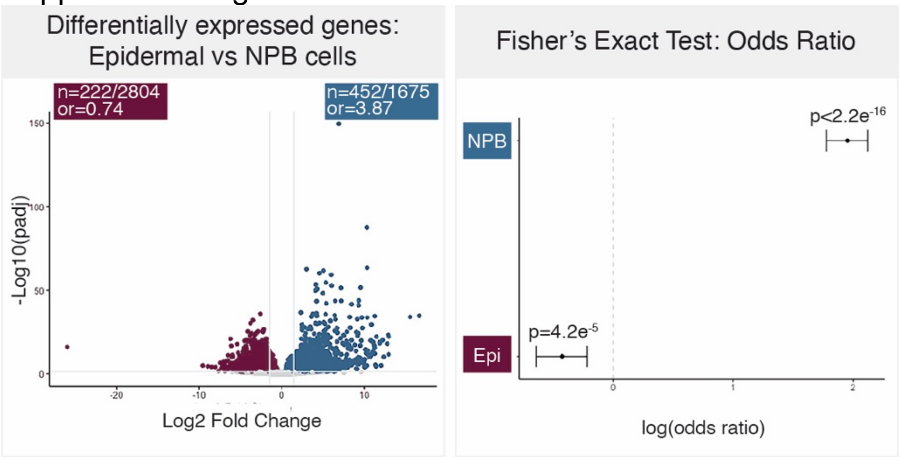


**Sup. Fig. 7. Fisher's exact tests on differentially expressed genes in relation to Sox3 binding.** Volcano plot showing differentially expressed genes between stage 13 epidermal and neural plate border-induced explants. Genes differentially expressed in epidermal cells are shown in maroon and in blue for neural plate border cells. Forrest plot displaying the log(odds ratio) for each group of differentially expressed genes (epidermal vs neural plate border) in relation to Sox3 binding in neural plate border-induced explants (St. 11.5).

Supplemental Figure 8

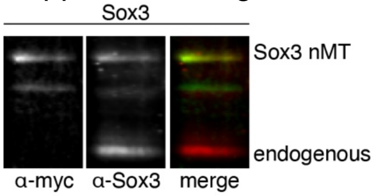


**Sup. Fig. 8. Sox3 was expressed at near endogenous levels for blastula (St. 9) ChIP-seq experiments.** Western blot for myc (green) and Sox3 (red) in embryos expressing *sox3* mRNA.

Supplemental Figure 9

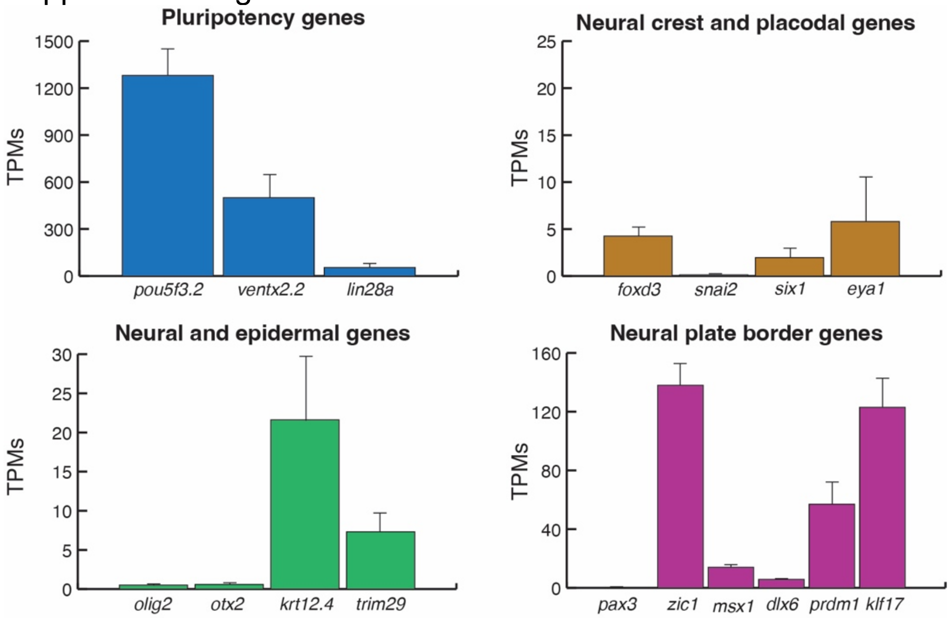


**Sup. Fig. 9. TPMs for genes in blastula stems cells.** TPMs for pluripotency genes (*pou5f3.2, ventx2.2* and *lin28a*), neural crest genes (*foxd3* and *snai2*), placodal genes (*six1* and *eya1*), neural genes (*olig2, otx2),* epidermal genes (*krt12.4, trim29*), and neural plate border genes (*pax3, zic1, msx1, dlx6, prdm1,* and *klf17*) in blastula stem cells (stage 9).

Supplemental Figure 10

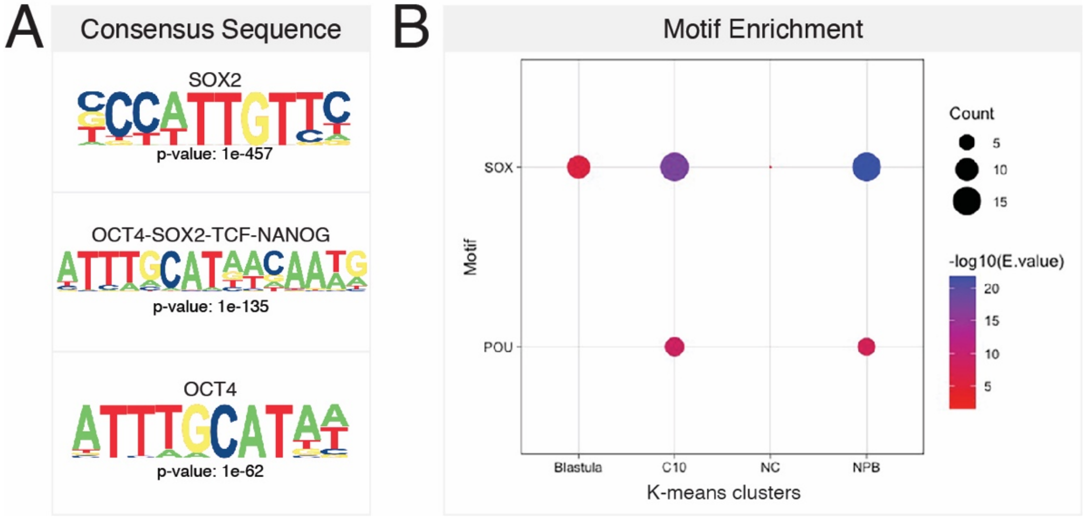


**Sup. Fig. 10. Motif analysis on shared Sox3 ChIP-seq peaks (St.9 and St. 11.5).** (A) HOMER motif consensus sequences and associated p-value. (B) Motif enrichment analysis, focusing on prevalence of SOX and POU motifs, on regions of Sox3 binding in the four k-means clusters.

Supplemental Figure 11

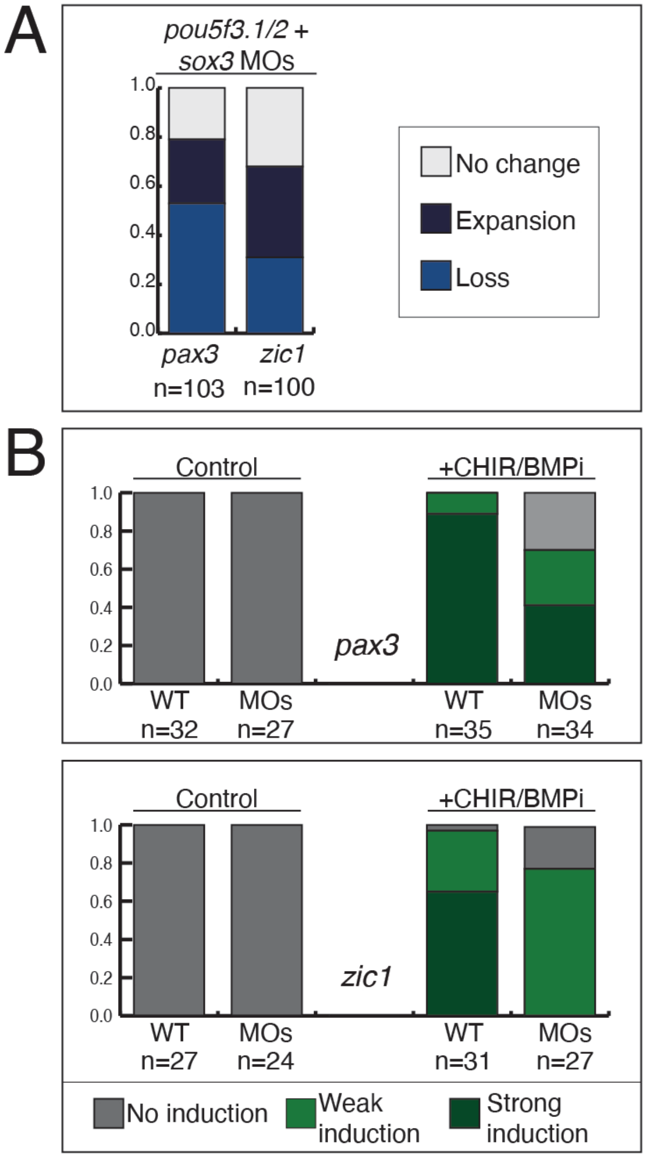


**Sup. Fig. 11. *Pou5f3.1/2 + sox3* triple morphant scoring.** (A) Stacked bar graphs with the percent of embryos with changes in gene expression (loss, expansion, no change) in *pou5f3.1+pou5f3.2+sox3* triple morphants (B) Stacked bar graphs with the percent of neural plate border-induced explants expressing *pax3* or *zic1*, indicating induction to a neural plate border state.
